# Ric8 proteins as the neomorphic partners of Gαo in *GNAO1* encephalopathies

**DOI:** 10.1101/2023.03.27.534359

**Authors:** Gonzalo P. Solis, Alexey Koval, Jana Valnohova, Mikhail Savitsky, Vladimir L. Katanaev

## Abstract

*GNAO1* mutated in pediatric encephalopathies encodes the major neuronal G-protein Gαo. Of >40 pathogenic mutations, most are single amino acid substitutions spreading across Gαo sequence. We perform extensive characterization of Gαo mutants showing abnormal GTP uptake and hydrolysis, and deficiencies to bind Gβγ and RGS19. Plasma membrane localization of Gαo is decreased for a subset of mutations that leads to epileptic manifestations. Pathogenic mutants massively gain interaction with Ric8A/B proteins, delocalizing them from cytoplasm to Golgi. Being general Gα-subunit chaperones and binding multiple other proteins, Ric8A/B likely mediate the disease dominance when engaging in neomorphic interactions with pathogenic Gαo. As the strength of Gαo-Ric8B interactions correlates with disease severity, our study further identifies an efficient biomarker and predictor for clinical manifestations in *GNAO1* encephalopathies.

**One-Sentence Summary:** Neomorphic mutations in Gαo gain dominant interactions with Ric8A/B, correlating with severity in pediatric encephalopathies.

## Introduction

Heterotrimeric G-proteins are the principal transducers of G protein-coupled receptors (GPCRs) – the biggest receptor family in animals – and consist of the α, β, and γ subunits. Sixteen human Gα-subunits fall into Gαi/o, Gαs, Gαq, and Gα12/13 subclasses. Upon activation, the cognate GPCR acts as a GEF (guanine-nucleotide exchange factor), catalyzing the GDP/GTP exchange on Gα and leading to the heterotrimer dissociation into Gα-GTP and Gβγ, both competent of downstream signaling (*1*). The intrinsic GTPase activity of Gα, further stimulated by RGS (regulator of G-protein signaling) proteins (*2*), leads to GTP hydrolysis and the resultant Gα-GDP can continue to signal reloading with GTP (*3*), or re-associates with Gβγ, closing the cycle (*1*).

Pediatric *GNAO1* encephalopathies are characterized by a spectrum of clinical manifestations including early onset epilepsy, motor dysfunctions, developmental delay, intellectual disability, and occasional brain atrophy (*4-7*). Caused primarily by dominant *de novo* mutations in *GNAO1*, the gene encoding the major neuronal G-protein Gαo, these encephalopathies lack efficient treatments. Missense mutations are most frequently seen in *GNAO1* encephalopathies and spread throughout the Gαo coding sequence, affecting conserved residues with critical functions in the G-protein (*8*). Thus far, pathological Gαo mutations have been described as loss-of-function (*9-11*), gain-of-function (*10, 12, 13*), or dominant-negative (*9-11, 14, 15*). This versatility of genetic manifestations has led us to propose that *GNAO1* encephalopathy mutations are neither of the above, but are instead of the neomorphic nature (*10*).

The neomorphic concept of *GNAO1* mutations imposes important constraints on the development of therapies (*10*), and further assumes that a novel mechanism is gained by pathologic Gαo mutants. Here, through a massive characterization of *GNAO1* mutations, we identify a uniform neomorphic feature: a strong gain-of-interaction, biochemical and cellular, with Ric8A and, more surprisingly, with Ric8B – the mandatory chaperones of all Gα-subunits (*16*). Furthermore, the neomorphic Gαo-Ric8B interaction emerges as a simple biomarker for the disease severity.

## Results

### Clinical assessment of *GNAO1* encephalopathy mutants

As representatives of pathogenic Gαo mutants, the following 16 were studied: G40R, G45E, S47G, D174G, L199P, G203R, R209C, C215Y, A227V, Y231C, Q233P, E237K, E246K, N270H, F275S, and I279N; a novel Q52R mutation (*17*) was also included in some analyses. The mutants were grouped following the OMIM catalog (https://omim.org/) that classifies *GNAO1* encephalopathy into two disorders with distinct clinical manifestations: “Developmental and Epileptic Encephalopathy-17” (DEE17; OMIM #615473) and “Neurodevelopmental Disorder with Involuntary Movements” (NEDIM; OMIM #617493) (table S1). While motor dysfunction is typically present in both DEE17 and NEDIM, the former additionally includes epilepsy. Another important category emanating from the clinical data is the disease onset, which we use as the clinical score for individual mutations (table S1). This analysis separates *GNAO1* encephalopathy cases into those with a *Very early onset* (<10 days postnatal; represented by G45E, L199P, F275S, and I279N), *Early onset* (≥10 days, <3 months postnatal; G40R, Q52R, D174G, G203R, A227V, Y231C, and N270H), *Late onset* (≥3 months, <2 years postnatal; S47G, R209C, E237K, and E246K), and *Very late onset* (≥2 years postnatal; C215Y and Q233P) (table S1). All *GNAO1* mutations leading to DEE17 (except for S47G) lay within the *Very early* and *Early onsets*, whereas all NEDIM-mutants are within the *Very late* and *Late onset*s. Correlating with the disease severity, the 29 DEE17-patients combined show a median disease onset of ∼43 days, as opposed to the ∼569 days of 31 NEDIM-patients (fig. S1A).

### Biochemical properties of *GNAO1* encephalopathy mutants

Gαo mutations affect residues within motifs controlling nucleotide binding and hydrolysis: P-loop, Switch regions I, II, and III, and other sites in the Ras-like domain (Fig. 1A). Thus, nucleotide uptake/hydrolysis is suspected to be aberrant across *GNAO1* mutations. Indeed, previous studies demonstrated that the Q52P/R mutants displayed complete loss of GTP uptake (*17*), R209H displayed a faster GTP uptake (*18*), while G203R, R209C, and E246K displayed faster GTP uptake and lost hydrolysis (*10*).

**Fig. 1.**
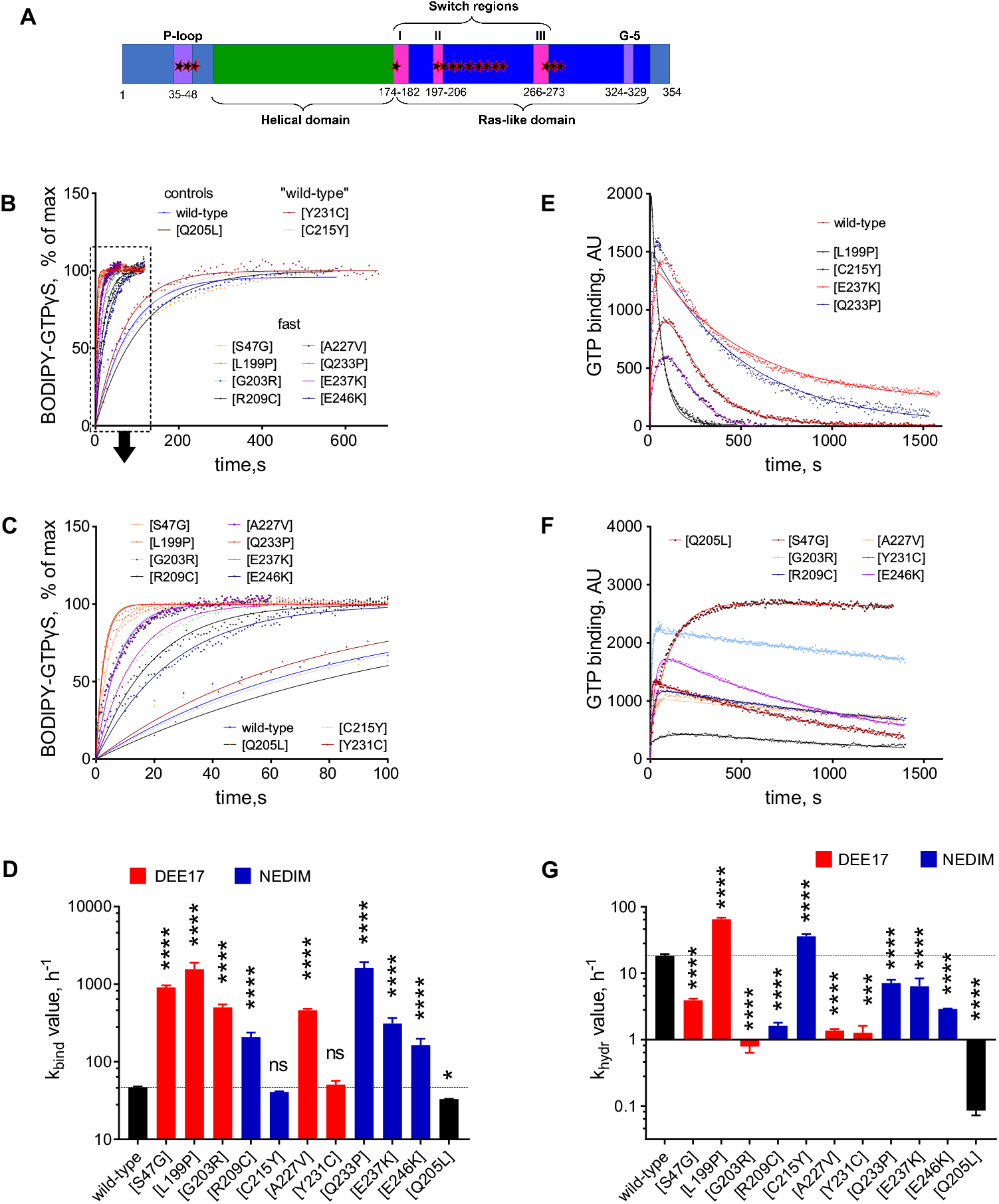
Spectrum of biochemical defects associated to *GNAO1* encephalopathy mutations. **(A)** Scheme of the mutated amino acid residues (stars) in the overall sequence Gαo. The residues are either located in the P-loop or in the Ras-like domain. **(B** and **C)** Representative curves of BODIPY-GTPγS binding to Gαo wild-type, encephalopathy mutants, and the GTPase-dead Q205L mutant (used as control). Most of Gαo mutants present strongly elevated binding rates, dotted-line box in **(B)** is expanded in **(C)**, whereas only two mutants (C215Y and Y231C) display near wild-type rates. **(D)** Quantification of the binding rate constant (k_bind_) of Gαo variants color-coded according to their association with the development of Developmental and Epileptic Encephalopathy-17 (DEE17; red bars) or Neurodevelopmental Disorder with Involuntary Movements (NEDIM; blue bars). **(E** and **F)** Representative curves of the course of BODIPY-GTP binding and hydrolysis by Gαo wild-type and active **(E)** or deficient/dead **(F)** mutants. **(G)** Quantification of the hydrolysis rate constant (k_hydr_). Note that data are adjusted to the plateau to highlight the differences in the binding rates **(B** and **C)**, while raw fluorescence units are shown in **(E** and **F)**, which are needed for the proper k_hydr_ calculation. Data in **(D)** and **(G)** are shown as means ± SD (*n*=3). ns is not significant, \**P* < 0.05 and \*\*\*\**P* < 0.0001 by one-way ANOVA followed by Dunnett’s multiple comparisons test.

We expressed in *E. coli* and affinity-purified the 16 pathologic Gαo mutants, along with Gαo wild-type and the classical GTPase-dead mutant Q205L as non-pathogenic controls (fig. S1B), and studied their GTP uptake/hydrolysis using BODIPY-GTPγS and BODIPY-GTP (*3, 10, 19, 20*). Six Gαo mutants (G40R, G45E, D174G, N270H, F275S, and I279N) were inactive, similarly to the Q52P/R mutants we studied earlier (*17*). All these mutations, including Q52P/R, lead to the more severe DEE17 disorder (fig. S1B and table S1). For the biochemically inactive mutants, the median disease onset of ∼31 days postnatal (13 patients) is, however, not significantly lower than the ∼52 days of the remaining 16 DEE17-patients (fig. S1C).

Analysis of the biochemically active variants reveals massive abnormalities in the GTP uptake and/or hydrolysis. First, the majority revealed dramatically faster rate of GTP uptake as compared to Gαo wild-type and Q205L (Fig. 1, B to D), generalizing the previous findings for G203R, R209C/H, and E246K (*10, 18*). Of the mutants studied, only C215Y and Y231C demonstrated near wild-type rates of GTP uptake; all the others increase the kinetic *k*_bind_ constant from 3.5-fold (E246K) to 34-fold (L199P) (Fig. 1D). Second, several Gαo mutants reveal strongly decreased rates of GTP hydrolysis (Fig 1, E to G), again like G203R, R209C, and E246K (*10*). Only two exceptions were seen: the kinetic *k*_hydr_ constant is increased for L199P (3.5-fold) and C215Y (2-fold), while the other mutants show a drop in *k*_hydr_ from 3-fold (E237K) to 14-fold (Y231C); the GTPase-dead Q205L drops this constant >200-fold (Fig 1G).

Given the strong increase in GTP uptake accompanied by a strong decrease in GTP hydrolysis, mutants are expected to be constitutively GTP-loaded. The only exception is C215Y, whose normal GTP uptake and enhanced GTP hydrolysis predict that this variant is preferably GDP-loaded compared to Gαo wild-type. To estimate the resulting preponderance of GTP charging, we performed simulations of the GDP/GTP cycling of the Gαo variants using the calculated *k*_bind_ and *k*_hydr_ (see Materials and Methods) (*21*). The ratio of the GTP-loaded protein to the GDP-bound is calculated as 2.56 to 1 for Gαo wild-type. This GTP/GDP ratio is strongly increased among most mutants, from 9.5-fold (L199P) to 245-fold (G203R) – as illustration, a 62.5-fold increase is calculated for Q205L. In contrast, C215Y shows a 2-fold decrease of the GTP/GDP ratio. We found, however, no significant correlation between disease onset and the GTP-loaded proportion of the mutants (fig. S1D).

### Cellular characterization of *GNAO1* encephalopathy mutants

Despite these insights into the biochemical properties of Gαo mutants, the complexity of cellular interactions, localizations, and signaling properties exceeds that of purified proteins. Thus, we moved next to massive cellular analyses, using Gαo variants with an internal GFP fusion allowing expression, localization, and co-immunoprecipitation (co-IP) analyses (*10, 17*). Transfection of the 16 pathologic mutants into the neuroblastoma N2a cell line demonstrates that most Gαo mutants have decreased expression compared to wild-type (fig. S2, A and B), with some (G40R, L199P, A227V, Y231C, N270H, and F275S) dropping to as low as ∼20% of wild-type. Noteworthy, the combined expression for the mutants related to DEE17, ∼35% of wild-type, was significantly lower than the ∼50% expression of the NEDIM-mutants (fig. S2C). In contrast, we found no correlation between disease onset and mutant expression levels (fig. S2D).

Previously, plasma membrane and Golgi localization of the G203R, R209C, and E246K mutants was observed (*10*), recapitulating the wild-type Gαo localization (*20*). In contrast, the Q52P/R mutants revealed severely decreased plasma membrane expression with maintained Golgi signal (*17*). Of note, Golgi localization – with or without plasma membrane binding – is indicative of a normal lipidation of Gαo (*22*). We systematically analyzed the localization pattern of the 16 Gαo mutants, revealing that they fall into two major groups. Both groups maintaining the Golgi localization, group 1 additionally maintains plasma membrane association, while group 2 strongly decreases it (Fig. 2, A to C and fig. S3A). A decrease in plasma membrane localization correlates significantly with a proportional increase in Golgi localization (fig. S3B). Combined, the plasma membrane and Golgi localizations of the mutants leading to DEE17 showed significant differences from their NEDIM counterparts (Fig. 2, D and E). We also find a significant correlation between disease onset and Gαo mutant localization at the plasma membrane, but not at Golgi (Fig. 2F and fig. S3C). These findings support our initial hypothesis that individual *GNAO1* mutations display DEE17 *vs*. NEDIM phenotypes depending on which of the two Gαo physiologic compartments – plasma membrane vs. Golgi – is primarily affected by the mutation (*23*).

**Fig. 2.**
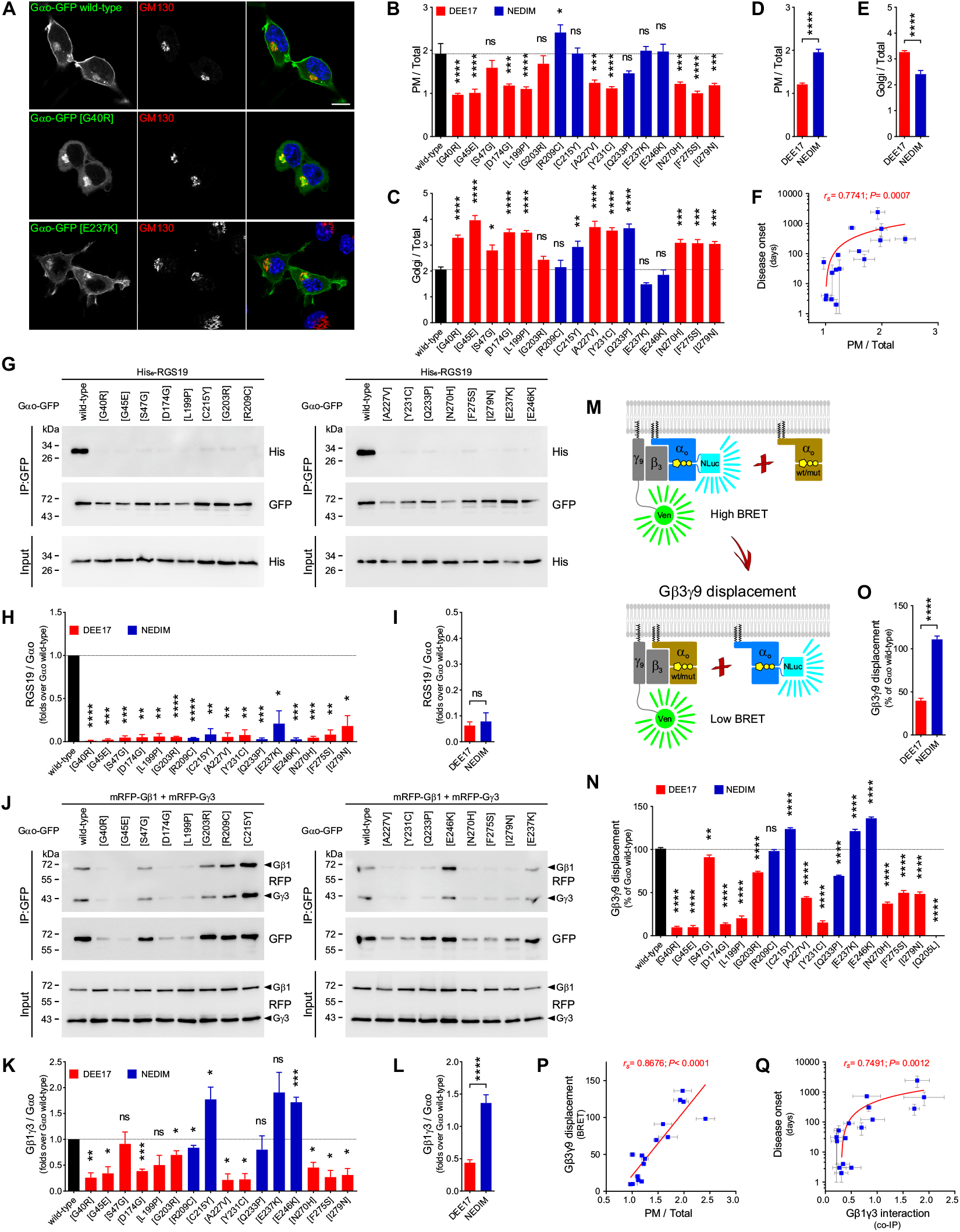
In-depth characterization of cellular properties of *GNAO1* mutations. **(A)** N2a cells expressing Gαo-GFP wild-type or the encephalopathy mutants G40R and E237K were immunostained against GM130 to visualize the Golgi apparatus and stained with DAPI in blue for nuclei. Scale bar, 10 µm. **(B** and **C)** Mean fluorescence intensity ratios of Gαo-GFP variants at the plasma membrane (PM; **B)** or Golgi **(C)** versus total cell (*n*=9-10). Bars are color-coded according to the involvement of Gαo mutants in the pathologies Developmental and Epileptic Encephalopathy-17 (DEE17; red) or Neurodevelopmental Disorder with Involuntary Movements (NEDIM; blue). **(D** and **E)** The combined localization at the PM **(D)** or Golgi **(E)** of the variants connected to DEE17 or NEDIM. **(F)** A scatterplot showing a significant positive correlation between Disease onset and PM localization of Gαo mutants. Note the log scale in the *y* axis. **(G)** N2a cells were co-transfected with His_6_-tagged RGS19 and Gαo-GFP variants, and immunoprecipitation (IP) was done with a nanobody against GFP. Coprecipitation of RGS19 was analyzed by Western blot (WB) using antibodies against GFP for Gαo and against His_6_-tag for RGS19. **(H** and **I)** Quantification of the co-IP of RGS19 by individual Gαo mutants (*n*=3) **(H)** and pooled in the DEE17 and NEDIM classes **(I). (J** to **L)** The interaction between Gαo-GFP wild-type and mutants with mRFP-Gβ1 and mRFP-Gγ3 was analyzed by IP and WB as in **(G)** using anti-GFP and anti-RFP antibodies **(J)**. Quantification of the Gαo-Gβ1γ3 interaction for individual Gαo variants (*n*=4-6) **(K)** and grouped in the DEE17 or NEDIM categories **(L). (M** to **O)** A scheme of the Gβ3γ9 displacement assay by BRET **(M)**. Wild-type Gαo internally tagged with NanoLuciferase (Gαo-NLuc) excites the cpVenus (Ven) fused to Gγ9 in the Gβ3γ9 heterodimer. The ability of non-tagged Gαo to displace Gβ3γ9 from Gαo-NLuc (reduction in the BRET signal) was quantified for Gαo wild-type, the encephalopathy mutants, and the GTPase-dead Q205L as control (*n*=4-9) **(N)**. The combined effect of the Gαo mutants on Gβ3γ9 displacement sorted in the DEE17 or NEDIM groups **(O). (P** and **Q)** Scatterplots illustrating a strong positive correlation between Gβ3γ9 displacement and PM localization **(P)**, and between disease onset and Gβ1γ3 co-IP **(Q)** of Gαo mutants. Note the log scale in the *y* axis of **(Q)**. All data is shown as means ± SEM. Data in **(B), (C)** and **(N)** were analyzed by one-way ANOVA followed by Dunnett’s multiple comparisons test, **(H)** and **(K)** by one-sample t-test, **(D), (E), (I), (L)**, and **(O)** by two-tailed Mann Whitney test, and **(F), (P)**, and **(Q)** by two-tailed Spearman correlation test; rank correlation coefficients (*r*_*s*_) and *P* values are indicated. ns is not significant, \**P* < 0.05, \*\**P* < 0.01, \*\*\**P* < 0.001, and \*\*\*\**P* < 0.0001.

### Interaction of *GNAO1* encephalopathy mutants with RGS19 and Gβγ

We previously showed that the interaction of some Gαo mutants (Q52P/R, G203R, R209C, and E246K) with RGS19, a major regulator of GTP hydrolysis on Gαo (*3*), is dramatically impaired (*10, 17*). We have now systematically analyzed RGS19-Gαo binding across the pathogenic mutants through co-IP using an anti-GFP nanobody (*10*). We see the loss of this interaction as an omnipresent phenomenon for *GNAO1* mutations, equally affecting mutants leading to DEE17 or NEDIM, and regardless of disease onset (Fig. 2, G to I and fig. S4A).

Our prior analyses of Gαo binding to Gβγ revealed different deviations from wild-type levels (*10, 17*). A systematic co-IP analysis of Gαo-GFP mutants co-expressed with mRFP-tagged Gβ1 and Gγ3 (*22*) uncovered varying perturbations of the Gβγ-Gαo mutant interaction (Fig. 2, J to L). As an independent confirmation of the co-IP studies, we employed a BRET displacement analysis (Fig. 2M). Specifically, we measured the ability of non-tagged Gαo (wild-type or mutants) to compete with the interaction between wild-type Gαo tagged with nanoLuciferase and Gβ3γ9 with a Venus-fusion (*10, 24*), revealing a perturbed Gβγ-Gαo pattern for the pathologic mutants similar to that seen in co-IPs (Fig. 2, N and O, and fig. S4B).

These findings permit making the firm conclusion that, as we suspected (*17*), decreased plasma membrane association strongly correlates with decreased interaction with Gβγ (Fig. 2P and fig. S4C). However, we still cannot determine which of the two is the cause, and which the consequence. Remarkably, a clear pattern emerges by both means to quantify Gαo-Gβγ interactions: mutations leading to DEE17 severely reduce the Gβγ binding, while NEDIM-mutations do not (Fig. 2, L and O).

When looking for possible correlations between the clinical score and the parameters analyzed so far, we found poor/non-existing correlations with Gαo expression levels (fig. S2D), Golgi localization (fig. S3C) or RGS19 interaction (fig. S4A). In contrast, a sizable correlation existed between disease onset and Gαo plasma membrane association (Fig. 2F) and Gβγ binding (Fig. 2Q and fig. S4D). However, as the range of pathologic Gβγ interactions goes from better than wild-type (E246K) to worse than wild-type Gαo (G40R), with the non-pathogenic Q205L displaying no interaction, the relative strength of the Gαo-Gβγ binding cannot serve as a simple biomarker to predict disease severity. Similarly, plasma membrane localization also cannot predict the clinical severity of the disease, as two mutations leading to DEE17 (S47G and G203R) showed near normal membrane localization (Fig. 2B and fig. S3A).

### Ric8A/B as neomorphic interaction partners of *GNAO1* encephalopathy mutants

The defective properties of pathogenic Gαo mutants that we have described so far do not explain the disease dominance, do not reveal a satisfactory biomarker to predict clinical severity, and do not identify the neomorphic function predicted by us (*10*). While searching for a neomorphic biomarker, we took into consideration the following observations: i) several recombinant Gαo mutants are biochemically inactive (fig. S1); ii) most mutants have decreased expression level (fig. S2); iii) despite constitutive GTP loading, the mutants fail to interact with RGS19 (Fig. 2, G and H). All these features hint at a potential folding problem, which prompted us to investigate Ric8A – GEF and chaperone of Gαo and other Gα-subunits (*16, 25*). In accordance with Ric8A chaperone functions that presume only transient interactions with its clients, we find very low capacity of GFP-tagged Ric8A to co-IP non-tagged Gαo wild-type and Q205L (Fig. 3, A and B). In contrast, all pathogenic Gαo mutants display a massive binding to Ric8A (Fig. 3, A to D). Note that the D174G mutants is poorly recognized by the anti-Gαo antibody used (E1; Fig. 3A), but its strong interaction with Ric8A was confirmed using another antibody (A2; Fig. 3C). Interestingly, the mutants leading to DEE17 combined show a slightly, but significantly, more pronounced Ric8A interaction than the NEDIM-mutants (Fig. 3E), although no significant correlation could be seen between disease onset and Ric8A binding (Fig. 3F).

**Fig. 3.**
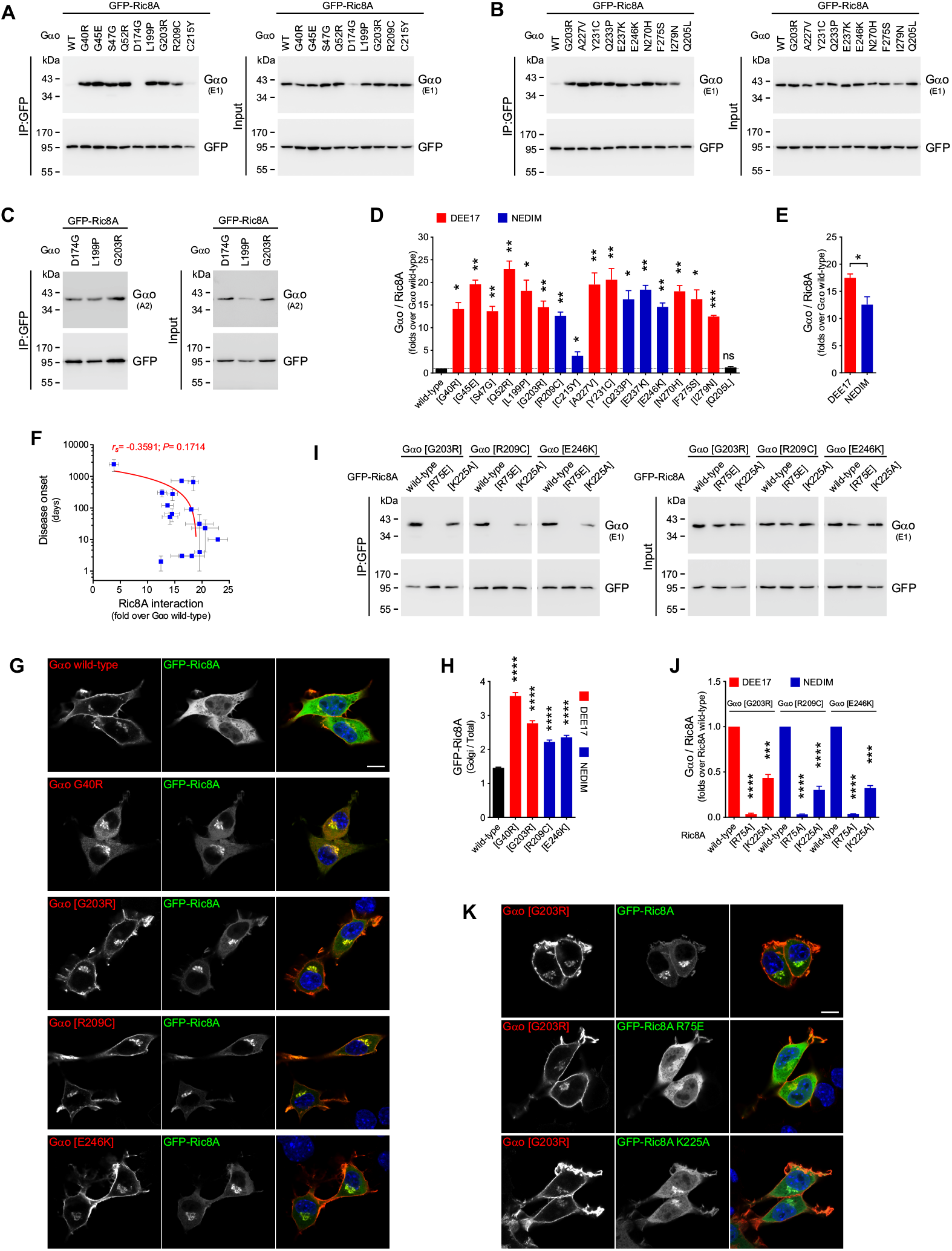
*GNAO1* mutants acquire a neomorphic interaction with Ric8A. **(A** to **C)** N2a cells were co-transfected with GFP-Ric8A and non-tagged Gαo wild-type, encephalopathy mutants, and the GTPase-dead Q205L mutant as control. The immunoprecipitation (IP) of GFP-Ric8A was achieved with a nanobody against GFP and the interaction with Gαo variants was determined by Western blot (WB), using antibodies against GFP and against Gαo clone E1 **(A** and **B)** or A2 **(C). (D)** Quantification of the co-IP of Gαo variants by Ric8A (*n*=3-4). Data are color-coded according to their involvement in Developmental and Epileptic Encephalopathy-17 (DEE17; red bars) or Neurodevelopmental Disorder with Involuntary Movements (NEDIM; blue bars). **(E)** The level of the Gαo-Ric8A interaction pooled according to DEE17 or NEDIM. **(F)** A scatterplot showing a non-significant negative correlation between Disease onset and Ric8A interaction of Gαo mutants. Note the log scale in the *y* axis. **(G)** N2a cells co-expressing GFP-Ric8A and non-tagged Gαo wild-type or selected encephalopathy mutants were immunostained against Gαo and stained with DAPI in blue for nuclei **(G)**. Scale bar, 10 µm. **(H)** Quantification of the mean fluorescence intensity ratio of GFP-Ric8A at the Golgi versus total cell (*n*=57-60). **(I)** N2a cells were co-transfected with GFP-Ric8A (wild-type or its chaperone-deficient mutants R75E and K225A) and non-tagged Gαo mutants (G203R, R209C or E246K). The IP of GFP-Ric8A constructs and immunodetection were done as in **(A). (J)** Quantification of the co-IP of Gαo mutants by the different Ric8A constructs (n=4-5). **(K)** N2a cells co-expressing Gαo G203R and GFP-Ric8A wild-type, R75E, or K225A were treated as in **(G)**. Scale bar, 10 µm. All data is shown as means ± SEM. Data in **(D)** and **(J)** were analyzed by one-sample t-test, **(E)** by two-tailed Mann Whitney test, **(F)** by two-tailed Spearman correlation test (rank correlation coefficient (*r*_*s*_) and *P* value are indicated), and **(H)** by one-way ANOVA followed by Dunnett’s multiple comparisons test. ns is not significant, \**P* < 0.05, \*\**P* < 0.01, \*\*\**P* < 0.001, and \*\*\*\**P* < 0.0001.

Ric8A is cytoplasmic, and this localization persists upon co-expression of Gαo wild-type or Q205L (Fig. 3, G and H, and fig. S5A). In contrast, every encephalopathy mutant induces a prominent neomorphic localization of GFP-Ric8A to the Golgi and, to a lower extent, plasma membrane (Fig. 3, G and H, and fig. S5A). Similar patterns were observed when HA-tagged Ric8A was co-transfected instead (fig. S6A). The extent of re-localization appears similar across the mutants, with the sole exception of C215Y with a weaker Golgi localization in accordance with its modest co-IP by Ric8A (Fig. 3, G and H, and fig. S5A). The Golgi-delocalization of Ric8A is a direct consequence (and not prerequisite) of its Gαo binding, as incubation with the N-myristoylation inhibitor DDD85646 (*22*), not preventing the co-IP of Gαo mutants by Ric8A (fig. S6, B and C), recovered the cytoplasmic localization of Ric8A (fig. S6, D to F). To assess the specificity of this neomorphic interaction, we introduced the point mutations into Ric8A known to abolish (R75E) or reduce (K225A) its chaperone activity (*26*). These Ric8A mutants showed a drastic decrease in binding the Gαo variants (Fig. 3, I and J), suggesting that pathogenic Gαo-Ric8A complexes are formed co-translationally. As expected, Ric8A constructs showed a clear reduction (K225A) or loss (R75E) of Golgi-delocalization by Gαo mutants (Fig. 3K and fig. S7, A and B).

We have previously described a *Drosophila* model of *GNAO1* encephalopathy that recapitulates clinical features seen in patients (*10*). The high degree of sequence identity and interchangeability of human and *Drosophila* Gαo (*8*) made us wonder whether the neomorphic Gαo-Ric8A interaction is evolutionary conserved in *Drosophila*. Indeed, we found that *Drosophila* Gαo G203R strongly interacts not only with *Drosophila* GFP-Ric8, but also with mammalian GFP-Ric8A (fig. S8A). Similarly, human Gαo G203R was efficiently co-precipitated by *Drosophila* Ric8 (fig. S8A). This result is somewhat surprising as, unlike the high degree of sequence identity between human and *Drosophila* Gαo reaching 84% (*8*), the identity between the sole *Drosophila* Ric8 ortholog and mammalian Ric8 is quite limited: 35% to Ric8A and 33% to Ric8B, with both mammalian Ric8 proteins being 47% identical (fig. S8B).

These findings prompted us to question whether the Gαo encephalopathy mutants can additionally gain a neomorphic interaction with Ric8B – the isoform ‘foreign’ for Gαo but specific instead for Gαs/olf (*16*). Surprisingly, we found that all Gαo mutants, except for C215Y, were significantly pulled-down by GFP-Ric8B, an effect not observed for Gαo wild-type and Q205L (Fig. 4, A to D). Remarkably, the DEE17-mutants combined show a much higher Ric8B binding than those leading to NEDIM (Fig. 4E), and a strong correlation is observed between disease onset and Ric8B interaction (Fig. 4F). We also observed that the mutants, but not Gαo wild-type, induce a cytoplasm-to-Golgi delocalization of Ric8B that also appears to follow the disease severity (Fig. 4G). Altogether, the strength of the neomorphic Ric8B binding provides the best predictive value for the clinical manifestations of *GNAO1* encephalopathy mutations.

**Fig. 4.**
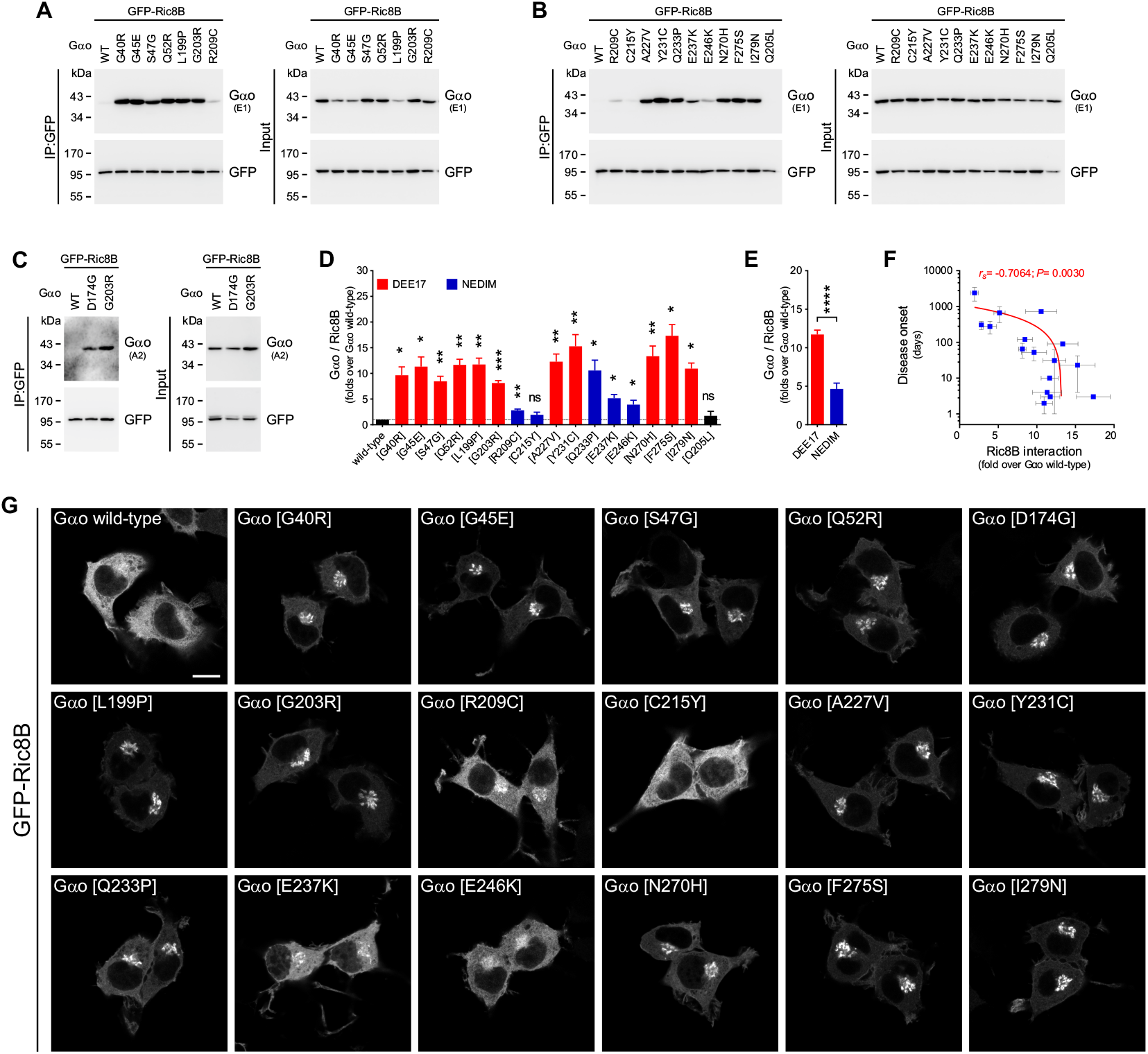
Ric8B neomorphic interaction with *GNAO1* mutants. **((A** to **C)** N2a cells were co-transfected with GFP-Ric8B and non-tagged Gαo wild-type, encephalopathy mutants, and Q205L as control. The immunoprecipitation (IP) of GFP-Ric8B was achieved with a nanobody against GFP and the interaction with Gαo variants was determined by Western blot (WB), using antibodies against GFP and against Gαo clone E1 **(A** and **B)** or A2 **(C). (D)** Quantification of Gαo pulled-down by Ric8B (*n*=3-5). Data are color-coded according to their involvement in Developmental and Epileptic Encephalopathy-17 (DEE17; red bars) or Neurodevelopmental Disorder with Involuntary Movements (NEDIM; blue bars). **(E)** Level of the Gαo-Ric8B neomorphic interactions according to DEE17 or NEDIM. **(F)** A scatterplot illustrating a significant negative correlation between Disease onset and the level of Gαo co-precipitated by Ric8B. Note the log scale in the *y* axis. **(G)** Representative images of the localization of GFP-Ric8B in N2a cells co-expressing Gαo wild-type or mutants (Gαo immunostaining not showed). Scale bar, 10 µm. All data is shown as means ± SEM. Data in **(D)** were analyzed by one-sample t-test, **(E)** by two-tailed Mann Whitney test, and **(F)** by two-tailed Spearman correlation test; rank correlation coefficient (*r*_*s*_) and *P* value are indicated. ns is not significant, \**P* < 0.05, \*\**P* < 0.01, \*\*\**P* < 0.001, and \*\*\*\**P* < 0.0001.

## Discussion

Recent advances in next-generation whole-exome/genome sequencing have uncovered a plethora of *GNAO1* mutations associated to a rare yet devastating pediatric encephalopathy. The number of patients steadily increases since the first reported cases in 2013 (*4*). The understanding of the molecular etiology underlying the pathology, however, remains unknown despite some current progress (*4, 9-11, 14, 15, 23*). In our previous study, we hypothesized that *GNAO1* encephalopathy mutations are neomorphic in nature (*10*), in the classical Muller categorization of genetic mutations. Citing from Muller’s seminal paper (*27*), neomorph represents a ‘*change in the nature of the gene at the original locus, giving an effect not produced, or at least not produced to an appreciable extent, by the original normal gene*’. In cancer, many oncogenic mutations in different genes, previously considered gain/loss-of-function, now emerge as neomorphs (*28*). Here, through a massive characterization of *GNAO1* mutations, we uncovered a neomorphic feature shared by all Gαo encephalopathy mutants: a strong gain-of-interaction with Ric8A and, even more surprisingly, with Ric8B.

Initially described as GEFs (*25*), Ric8 proteins have subsequently emerged as mandatory chaperones for Gα-subunits (*16*), with Ric8A responsible for the Gαi/o, Gαq, and Gα12/13 subfamilies, and Ric8B solely for Gαs/olf. Ric8 interactions with Gα subunits are highly specific, with Ric8B been unable to engage members of the Gαi/o class, Gαo included (*25, 29*). As the main determinant for Ric8B specificity resides in the extended C-terminal α5 helix of Gαs/olf, shorter in Gαi/o members (*30*), it is tempting to speculate that *GNAO1* mutations alter Gαo structure in ways that differentially accommodate its α5 helix to Ric8B. Given the obligate chaperoning function of Ric8A/B for all Gα-subunits, the neomorphic Gαo-Ric8 interaction seen for the pathogenic variants may lie at the core of the disease dominance affecting not only Gαo signaling, but also disbalancing the entire neuronal GPCR signaling networks. Ric8A and Ric8B also interact with a number of other partners with divergent activities (BioGRID #121946 and #120486; https://thebiogrid.org/); these interactions might also be affected by the neomorphic Gαo-Ric8A/B binding and may represent alternative/additional means of disease dominance in *GNAO1* encephalopathies. Noteworthy, the neomorphic Gαo-Ric8A/B interaction is an attractive target for future drug discovery efforts.

Our study provides the first systematic investigation of a large panel of pathogenic Gαo mutants across several biochemical and cellular assays, revealing that the mutants tend to bind GTP faster but lose GTP-hydrolysis, lose interaction to RGS19, display varying deficiencies in plasma membrane expression and Gβγ binding, and gain a neormophic Ric8A/B interaction. However, Gαo C215Y stands out the group as it shows an increased Gβγ interaction, normal subcellular localization, and a moderate binding exclusively to Ric8A but not Ric8B. C215Y also loses RGS19 binding, although this might reflect its nucleotide-state in cells, as it is the only Gαo mutant predicted to have a higher GDP-loading than wild-type. Thus, the C215Y variant, as the T241_N242insPQ (c.724-8G>A) mutant recently characterized by us (*31*), might fall into a separate category of mild *GNAO1* encephalopathy with adolescent or adult-onset, as suggested (*32*).

Out of multiple deficiencies of pathogenic Gαo variants, the interaction with Ric8B emerges not only as the most unexpected neomorphic feature of the disease, but also as the best predictor of its severity. *GNAO1* encephalopathy is a recently discovered disorder, and most known patients are infants with unclear prognosis. The molecular outcome of many *GNAO1* mutations remains unknown, as sequencing continues identifying more patients, often with novel mutations. With this background, a simple biomarker of the disease severity is in a high demand. Our work discovers the Gαo-Ric8B interaction as such a biomarker, with the strength of the interaction correlating with disease severity. A simple test measuring Ric8B interaction with the new variant may become the routine way for medical geneticists and pediatric neurologists to evaluate the expected disease severity, long-term prognosis, and treatment.

In sum, our work sheds light on the molecular etiology of *GNAO1* encephalopathy, identifies the neomorphic intermediates of the disease dominance, provides a much-needed biomarker for assessment of disease severity, and might pave the way for future drug discovery towards this disorder.

## Acknowledgments

We thank Dr. Mikhail Kryuchkov for critical discussion of this work, S. Troccaz for the technical assistance, and the members of the Bioimaging core facility of the CMU for assistance in microscopy.

## Author contributions

M.S. performed molecular cloning. A.K. performed biochemical experiments and BRET analysis. G.P.S. and J.V. performed cellular experiments. G.P.S and V.L.K. designed and supervised the study and wrote the manuscript. All authors contributed to revising the manuscript.

## References

1. W. M. Oldham, H. E. Hamm, Heterotrimeric G protein activation by G-protein-coupled receptors. Nat Rev Mol Cell Biol 9, 60–71 (2008).

2. E. M. Ross, T. M. Wilkie, GTPase-activating proteins for heterotrimeric G proteins: regulators of G protein signaling (RGS) and RGS-like proteins. Annu Rev Biochem 69, 795–827 (2000).

3. C. Lin et al., Double suppression of the Galpha protein activity by RGS proteins. Molecular cell 53, 663–671 (2014).

4. K. Nakamura et al., De Novo mutations in GNAO1, encoding a Galphao subunit of heterotrimeric G proteins, cause epileptic encephalopathy. Am J Hum Genet 93, 496–505 (2013).

5. T. Schirinzi et al., Phenomenology and clinical course of movement disorder in GNAO1 variants: Results from an analytical review. Parkinsonism Relat Disord 61, 19–25 (2019).

6. M. Kelly et al., Spectrum of neurodevelopmental disease associated with the GNAO1 guanosine triphosphate-binding region. Epilepsia 60, 406–418 (2019).

7. E. Axeen et al., Results of the First GNAO1-Related Neurodevelopmental Disorders Caregiver Survey. Pediatr Neurol 121, 28–32 (2021).

8. M. Savitsky, G. P. Solis, M. Kryuchkov, V. L. Katanaev, Humanization of Drosophila Gαo to Model GNAO1 Paediatric Encephalopathies. Biomedicines 8, 395 (2020).

9. B. S. Muntean et al., Gαo is a major determinant of cAMP signaling in the pathophysiology of movement disorders. Cell Rep 34, 108718 (2021).

10. Y. A. Larasati et al., Restoration of the GTPase activity and cellular interactions of Galpha(o) mutants by Zn(2+) in GNAO1 encephalopathy models. Sci Adv 8, eabn9350 (2022).

11. D. Wang et al., Genetic modeling of GNAO1 disorder delineates mechanisms of Gαo dysfunction. Hum Mol Genet 31, 510–522 (2022).

12. H. Feng et al., Movement disorder in GNAO1 encephalopathy associated with gain-of-function mutations. Neurology 89, 762–770 (2017).

13. L. Song et al., Identification of functional cooperative mutations of GNAO1 in human acute lymphoblastic leukemia. Blood 137, 1181–1191 (2021).

14. M. Di Rocco et al., Caenorhabditis elegans provides an efficient drug screening platform for GNAO1-related disorders and highlights the potential role of caffeine in controlling dyskinesia. Hum Mol Genet 31, 929–941 (2022).

15. D. Silachev et al., Mouse models characterize GNAO1 encephalopathy as a neurodevelopmental disorder leading to motor anomalies: from a severe G203R to a milder C215Y mutation. Acta Neuropathol Commun 10, 9 (2022).

16. M. Gabay et al., Ric-8 proteins are molecular chaperones that direct nascent G protein α subunit membrane association. Sci Signal 4, ra79–ra79 (2011).

17. G. P. Solis et al., Pediatric Encephalopathy: Clinical, Biochemical and Cellular Insights into the Role of Gln52 of GNAO1 and GNAI1 for the Dominant Disease. Cells 10, 2749 (2021).

18. C. L. Larrivee et al., Mice with GNAO1 R209H Movement Disorder Variant Display Hyperlocomotion Alleviated by Risperidone. J Pharmacol Exp Ther 373, 24–33 (2020).

19. D. Kopein, V. L. Katanaev, Drosophila GoLoco-protein pins is a target of Galpha(o)-mediated G protein-coupled receptor signaling. Mol Biol Cell 20, 3865–3877 (2009).

20. G. P. Solis et al., Golgi-Resident Galphao Promotes Protrusive Membrane Dynamics. Cell 170, 939–955 (2017).

21. V. L. Katanaev, M. Chornomorets, Kinetic diversity in G-protein-coupled receptor signalling. Biochem J 401, 485–495 (2007).

22. G. P. Solis et al., Local and substrate-specific S-palmitoylation determines subcellular localization of Gαo. Nat Commun 13, 2072 (2022).

23. G. P. Solis, V. L. Katanaev, Galphao (GNAO1) encephalopathies: plasma membrane vs. Golgi functions. Oncotarget 9, 23846–23847 (2018).

24. H. Schihada, R. Shekhani, G. Schulte, Quantitative assessment of constitutive G protein-coupled receptor activity with BRET-based G protein biosensors. Sci Signal 14, eabf1653 (2021).

25. G. G. Tall, A. M. Krumins, A. G. Gilman, Mammalian Ric-8A (synembryn) is a heterotrimeric Galpha protein guanine nucleotide exchange factor. Journal of Biological Chemistry 278, 8356–8362 (2003).

26. A. B. Seven et al., Structures of Gα Proteins in Complex with Their Chaperone Reveal Quality Control Mechanisms. Cell Rep 30, 3699–3709.e3696 (2020).

27. H. J. Muller, Further studies on the nature and causes of gene mutations. Proceedings of the Sixth International Congress of Genetics, Ithaca, New York. 1, 213–255 (1932).

28. V. Takiar, C. K. Ip, M. Gao, G. B. Mills, L. W. Cheung, Neomorphic mutations create therapeutic challenges in cancer. Oncogene 36, 1607–1618 (2017).

29. P. Chan, M. Gabay, F. A. Wright, G. G. Tall, Ric-8B is a GTP-dependent G protein alphas guanine nucleotide exchange factor. J Biol Chem 286, 19932–19942 (2011).

30. M. M. Papasergi-Scott et al., Structures of Ric-8B in complex with Galpha protein folding clients reveal isoform specificity mechanisms. Structure, (2023).

31. A. Koval et al., In-depth molecular profiling of an intronic GNAO1 mutant as the basis for personalized high-throughput drug screening. Med 4, in press (2023).

32. T. Wirth et al., Highlighting the Dystonic Phenotype Related to GNAO1. Mov Disord 37, 1547–1554 (2022).

