## Supplementary material for "Ric8 proteins as the neomorphic partners of Gαo in *GNAO1* encephalopathies": Material & Methods, Table S1-S2, Fig. S1-S8

### Materials and Methods

#### Antibodies and reagents

Primary antibodies (Abs) for immunofluorescence (IF) and Western blots (WBs): monoclonal Abs (mAbs) anti-Gαo-A2 (sc-13532; WB: 1/50, IF: 1/50), anti-Gαo-E1 (sc-393874; WB: 1/250), and anti-mRFP/DsRed2 (sc-101526; WB: 1/250) were from Santa Cruz Biotechnology, mAb anti-$His_6$ (34650; WB: 1/1000) from Qiagen, mAb anti-GM130 (610823; IF: 1/500) from BD Biosciences, and mAb anti-α-tubulin (T6199; WB: 1/2000) from Sigma-Aldrich. Polyclonal antibody (pAb) anti-GFP (GTX113617; WB: 1/2000) was from GeneTex and pAb against HA-tag (ab9110) was from Abcam. The pAb against *Drosophila* Gαo (WB: 1/2000) was previously published (*33*). All secondary Abs for immunofluorescence (IF) and Western blots (WBs) were from Jackson ImmunoResearch: anti-Mouse Cy3-conjugated (115-165-146; IF: 1/500), anti-Rabbit Alexa488-conjugated (111-545-144; IF: 1/500), anti-Mouse Horseradish peroxidase (HRP)-conjugated (115-035-146; WB: 1/5000), and anti-Rabbit HRP-conjugated (111-035-144; WB: 1/5000). DAPI (32670) was from Sigma-Aldrich, VECTASHIELD Mounting Medium (H-1400) from Vector Laboratories, Glutathione Sepharose 4B beads (17075601) from Cytiva, and DDD85646 (13839) from Cayman Chemical.

#### Plasmids and molecular cloning

The plasmids encoding for non-tagged Gαo, $His_6$-tagged Gαo and Gαo-GFP (GFP insertion between residues G92 and I93) for the wild-type and mutants (Q52R, G203R, Q205L, R209C, and E246K), $His_6$-RGS19, mRFP-Gβ1, mRFP-Gγ3, and non-tagged *Drosophila* Gαo were previously described (*3, 10, 19, 34, 35*). Additional Gαo mutants G40R, G45E, S47G, D174G, L199P, C215Y, A227V, Y231C, Q233P, E237K, N270H, F275S, and I279N, and *Drosophila* Gαo G203R were obtained by point mutagenesis using the oligonucleotide primers (all primers used in this study are listed in table S2). To generate GFP-Ric8A, the Ric8A sequence was amplified by PCR from the HA-Ric8A plasmid (*36*) (kindly provided by Yijuang Chern; Academia Sinica, Taiwan) and cloned in frame into the SalI/PspOMI sites of pEGFP-C3 (Clontech). The R75E and K225A mutations were introduced into GFP-Ric8A by point mutagenesis. Ric8B was PCR-amplified from pB-Ric8B (*37*) (Addgene plasmid #129457) and cloned in frame into the XhoI/EcoRI sites of pEGFP-C1 (Clontech), producing the GFP-Ric8B construct. To clone a GFP-fusion of dRic8 (*Drosophila*), the dRic8 sequence was PCR-amplified from the pMT-GFP-dRic8 (*38*) (provided by Stephen L. Rogers; University of North Carolina at Chapel Hill) and inserted in frame into the BsrGI/PspOMI sites of pEGFP-C1.

#### Recombinant protein purification

$His_6$-tagged Gαo proteins were expressed in *Escherichia coli* Rosettagami (Novagen, 71351) as previously described (*10*). Briefly, transformed bacteria were grown in baffled flasks at 37°C to an $OD_{600}$ of 0.6, cooled down to 18°C for at least 30 min before induction with 1 mM Isopropyl-β-D-thiogalactopyranoside, and were additionally grown overnight at 18°C. Bacteria were harvested by centrifugation 3,500xg at 4°C and resuspended in TBS (20 mM Tris-HCl (pH 7.5) and 150 mM NaCl) supplemented with 1 mM PMSF and 30 mM imidazole (all Sigma-Aldrich). Cells were disrupted with a OneShot high-pressure cell press disruptor

(CONSTANT Systems) at 0.7Kbar and extracts were cleared by centrifugation at 15,000xg for 15 min at 4°C. Supernatants were incubated with Ni-NTA Agarose beads (QIAGEN) overnight in a rotary shaker at 4°C. Beads were washed twice with 10 volumes of wash buffer (TBS supplemented with 10 mM imidazole) and bound proteins were GDP-loaded in TBS supplemented with 3% glycerol, 10 mM $MgCl_2$, 0.1 mM DTT, and 200 μM GDP. Beads were washed two more times with at least 10 volumes of ice-cold wash buffer and finally eluted with TBS containing 300 mM imidazole. Imidazole was removed by buffer exchange to TBS using Vivaspin Centrifugal concentrators. Protein concentration was measured using the Bradford assay, and the purity was analyzed using SDS-PAGE followed by Coomassie staining.

**GTP-binding and hydrolysis assay**

The GTP-binding and hydrolysis assay using BODIPY-GTP or BODIPY-GTPγS (both from Invitrogen) was performed as described (*3*). Shortly, $His_6$-Gαo recombinant proteins were diluted to 1 μM in reaction buffer (TBS supplemented with 10 mM $MgCl_2$ and a 0.5% BSA). The mixture was then pipetted into black 384-well plates (Greiner), and BODIPY-GTP or BODIPY-GTPγS was added into the wells to 1 μM final concentration. Fluorescence measurements were performed at 28°C in a Tecan Infinite M200 PRO plate reader with excitation at 485 nm and emission at 530 nm. In all cases the fluorescent ligand was injected into the wells as half of the final volume of the reaction mixture using the injector system of the plate reader. The GTP-binding and hydrolysis data of Gαo wild-type were fit to obtain the $k_{bind}$ and $k_{hydr}$ rate constants as previously described (*3*), setting the end point as the baseline. For Gαo mutants with strongly impaired GTP hydrolysis, the maximal duration of the assay was not sufficient to reach complete hydrolysis and thus the $k_{hydr}$ was extrapolated using the available BODIPY-GTP curve with the initial fluorescence value as a projected baseline.

**Modeling Gαo$^{GTP}$/Gαo$^{GDP}$ ratios.**

Considering a ca. 10-fold excess of free GTP to GDP in cells, setting the cellular concentration of Gαo at 10 μM, and considering a ca. 100-fold excess of guanine nucleotides over the G proteins (*21*), in the absence of GEFs, GAPs, or GDIs the ratio of the GTP-loaded Gαo to that in the GDP-bound state is determined by the simple reactions:

$$\frac{[\text{G}\alpha\text{o}^{GDP}]}{dt} = k_{hydr}[\text{G}\alpha\text{o}^{GTP}] - k_{bind}[\text{G}\alpha\text{o}^{GDP}];$$

$$\frac{[\text{G}\alpha\text{o}^{GTP}]}{dt} = k_{bind}[\text{G}\alpha\text{o}^{GDP}] - k_{hydr}[\text{G}\alpha\text{o}^{GTP}]$$

Knowing the $k_{bind}$ and $k_{hydr}$ values for wild-type and mutant Gαo, these differential equations were numerically solved with using the PLAS (Power Law Analysis and Simulation) software (https://github.com/SMRUCC/PLAS.NET) as described (*21*). Resulting equilibrium concentrations of Gαo$^{GTP}$ and Gαo$^{GDP}$ provided the [Gαo$^{GTP}$] / [Gαo$^{GDP}$] ratios, shown in fig. S1D.

**Cell lines and culture conditions**

The mouse neuroblastoma Neuro-2a (N2a; ATCC, CCL-131) were maintained in MEM (Thermo Fisher Scientific), supplemented with 10% FCS, 2 mM L-glutamine, 1 mM pyruvate, and 1% penicillin-streptomycin at 37°C and 5% CO2. Human HEK293T (ATCC, CRL-3216) cells grew in DMEM (Thermo Fisher Scientific), supplemented as above and under the same culture conditions. All vector transfections were carried out with X-tremeGENE HP (Roche, XTGHP-RO) or FuGENE HD (Promega, E2311) according to manufacturer's instructions.

**Immunofluorescence and microscopy**

N2a cells were seeded on culture plates ($1x10^5$ cells/well), 24 h later were transfected for 7 h, trypsinized and seeded on poly-L-lysine-coated coverslips in complete MEM for additional 15-17 h before fixation. When indicated, cells were seeded in complete MEM supplemented with 10 µM DDD85646 or DMSO as control. Cells were fixed with 4% paraformaldehyde in PBS for 20 min, permeabilized for 1 min using ice-cold PBS supplemented with 0.1% Triton X-100, and blocked for 30 min with PBS supplemented with 1% BSA. Cells were then incubated with the primary Abs in blocking buffer for 2 h at room temperature (RT), washed, incubated with secondary Abs and DAPI also in blocking buffer for 2 h at RT, and coverslips were finally mounted with VECTASHIELD on microscope slides. Cells were recorded with a Plan-Apochromat 63x/1.4 oil objective on a LSM800 Confocal Microscope using the ZEN 2.3 software (all Zeiss). When required, mean fluorescence intensity was determined from confocal images using ImageJ v1.53t (National Institutes of Health). Images were not recorded using the same confocal settings, therefore ratio fluorescence values, such as Golgi fluorescence vs. total fluorescence or plasma membrane (PM) vs. total, were used for quantifications as previously validated (*39*). All images were finally edited using ZEN lite 3.3 (Zeiss) and CorelDRAW 2020 (Corel).

**PM and Golgi localization**

N2a cells were transfected with 0.5 µg of Gαo-GFP wild-type or mutant plasmids, immunostained against GM130 to visualize the Golgi apparatus and DAPI for Nuclei (as indicated above). For Ric8A/B localization studies, N2a cells were co-transfected with non-tagged Gαo (0.2 µg) and GFP-Ric8A, HA-Ric8A or GFP-Ric8B (0.4 µg), immunostained against Gαo and when needed HA-tag, and stained with DAPI (not shown for all images). To avoid the interference due to cell variability of expression of the constructs, mean fluorescence intensity was measured at the Golgi region as well as at the total cell area, and ratio values were used to determine their relative Golgi accumulation. Likewise, mean fluorescence intensity was determined at an unbroken PM region without membrane protrusions, and the ratio over total cell fluorescence was used to define relative PM content for each Gαo-GFP construct.

**Biochemical analyses**

For Gαo expression analysis, N2a cells were seeded on culture plates ($1x10^5$ cells/well) and 24 h later were transfected with 0.5 µg of Gαo-GFP wild-type or mutants. After additional 24 h, cells were harvested with

Accutase (Thermo Fisher Scientific), lysed in Laemmli buffer with sonication, boiled at 95°C for 5 min and finally analyzed by SDS-PAGE and Western blots using antibodies against GFP and α-tubulin as loading control. HRP-conjugated secondary antibodies were used for enhanced chemiluminescence (ECL) detection in a Fusion FX6 Edge system (Vilber). Quantification of all blots was done using ImageJ v1.53c, and images were edited using EvolutionCapt v18.11 (Vilber) and CorelDRAW 2020.

**Co-Immunoprecipitations**

The recombinant GST-tagged Nanobody against GFP (*40*) expressed in *Escherichia coli* Rosettagami was purified with Glutathione Sepharose 4B beads according to manufacturer's instructions. Protein purity was assessed by SDS-PAGE and Coomassie blue staining.

N2a cells were seeded on plates (2x$10^5$ cells /well) and cultured for 48h before co-transfection with 3 µg total DNA using the following combinations: Gαo-GFP and mRFP-Gβ1/Gγ3 (1 µg each), Gαo-GFP and $His_6$-RGS19 (1.5 µg each), or GFP-Ric8A/B and non-tagged Gαo (1.5 µg each). When indicated, DDD85646 was added to a 10 µM final concentration 7 h after transfection (DMSO was used as control). After 24 h transfection, cells were resuspended with ice-cold GST-lysis buffer (20 mM Tris-HCl, pH 8.0, 1% Triton X-100 and 10% glycerol in PBS) supplemented with a protease inhibitor cocktail (Roche) and passed >10 times through a 25G needle. Extracts were cleared by centrifugation at 15,000xg for 15 min at 4°C, and supernatants were incubated with 2 µg of purified GST-tagged GFP-Nanobody for 30 min on ice. Then, 20 µL of Glutathione Sepharose 4B beads were added, samples were rotated overnight at 4°C, beads were repeatedly washed with GST-lysis buffer, prepared for SDS-PAGE, and finally analyzed by Western blot using antibodies against GFP, mRFP, $His_6$-tag, and/or Gαo, followed by incubation with HRP-conjugated secondary antibodies for ECL detection as mentioned above. If not quantified, co-immunoprecipitations were done in duplicate with very similar outcomes.

**Gβ3γ9 displacement assay by BRET**

The plasmid Go1-CASE encoding for nanoLuciferase-tagged Gαo, Gβ3, and Venus-tagged Gγ9 was kindly supplied by Gunnar Schulte (*24*) (Karolinska Institutet, Sweden). HEK293T cells were co-transfected with the Go1-CASE plasmid and non-tagged Gαo wild-type or mutants at a 1:1 ratio. Twelve hours after transfection, cells were resuspended in complete DMEM and seeded in transparent-bottom black 384-well plates (6,000 cells/well). After 24 hours, the medium was replaced by 10 µl of PBS, and Furimazine was injected in an equal volume of PBS to a 10 µM final concentration immediately before measurement. GFP and Nanoluc signals were read at intervals of approximately 1.6s using built-in filter NanoBRET filter system for ~30s, and ratios averaged.

**Multiple sequence alignment**

The multiple sequence alignment for Ric8 proteins was done using the Clustal Omega tool of EMBL-EBI (*41*) and edited using the Jalview 2.11.2.6 software (*42*). The following sequences were used: Ric8A *Mus musculus* (NP_444424.1), Ric8B *Mus musculus* (NP_898995.1) and dRic8 *Drosophila melanogaster* (NP_001285048.1).

**Table S1. Clinical manifestations of the *GNAO1* encephalopathy mutations analyzed**. Online Mendelian Inheritance in Man (OMIM) entries for *GNAO1* encephalopathy: Developmental and Epileptic Encephalopathy-17 (**DEE17**; #615473) and Neurodevelopmental Disorder with Involuntary Movements (**NEDIM**; #617493).

| Amino acid change | Nucleotide change | # of patients described | Age of onset | Epilepsy | Movement disorder (excl. hypotonia) | Developmental delay | Brain alterations (MRI) | Ref. | OMIM category and score (days of onset, mean ± sem) |
|---|---|---|---|---|---|---|---|---|---|
| G40R | c.118G>C | 5 | 2.5m | yes | yes | yes | yes | (*6*) | DEE17 |
| | c.118G>A | | birth | yes | no | yes | no | (*43*) | 52 ± 22 (n=5): Early |
| | c.118G>C | | 2m | yes | yes | yes | yes | (*44*) | |
| | c.118G>A | | 6d | yes | no | N/A | yes | (*45*) | |
| | c.118G>C | | 4m | yes | no | yes | yes | (*46*) | |
| G45E | c.134G>A | 1 | 4d | yes | yes | yes | yes | (*47*) | DEE17<br>4 (n=1): Very early |
| S47G | c.139A>G | 1 | 4m | yes | yes | yes | yes | (*44*) | DEE17<br>120 (n=1): Late |
| Q52R | c.155A>G | 1 | 1.5w | yes | yes | yes | yes | (*17*) | DEE17<br>10 (n=1): Early |
| D174G | c.521A>G | 1 | 29d | yes | no | yes | yes | (*4*) | DEE17<br>29 (n=1): Early |
| L199P | c.596T>C | 1 | 3d | yes | yes | yes | yes | (*48*) | DEE17<br>3 (n=1): Very early |
| G203R | c.607G>A | 9 | 7m | yes | yes | yes | yes | (*4*) | DEE17 |
| | | | 1d | yes | yes | yes | yes | (*5*) | 65 ± 29 (n=9): Early |
| | | | 12d | yes | yes | yes | yes | (*5*) | |
| | | | 7m | yes | yes | yes | yes | (*49*) | |
| | | | 7d | yes | yes | yes | yes | (*50*) | |
| | | | 1m | yes | yes | yes | yes | (*51*) | |
| | | | 3m | yes | yes | yes | yes | (*52*) | |
| | | | 9d | yes | yes | yes | yes | (*52*) | |
| | c.607G>A | | 12d | yes | yes | yes | no | (*46*) | |
| R209C | c.625C>T | 13 | 6m | no | yes | yes | yes | (*53*) | NEDIM |
| | c.626G>A | | 2y | no | yes | yes | no | (*54*) | 305 ± 81 (n=10): Late |
| | c.625C>T | | infancy | no | yes | yes | no | (*55*) | |
| | c.625C>T | | 6m | yes | yes | yes | no | (*56*) | |
| | c.625C>T | | 3-4m | no | yes | yes | no | (*57*) | |
| | c.625C>T | | 2y | yes | yes | yes | N/A | (*58*) | |
| | c.625C>T | | N/A | yes | yes | yes | yes | (*59*) | |
| | c.625C>T | | infancy | yes | yes | yes | yes | (*60*) | |
| | c.625C>T | | infancy | yes | yes | yes | yes | (*60*) | |
| | c.625C>T | | 7m | yes | yes | yes | yes | (*50*) | |
| | c.625C>T | | 1.5y | no | yes | no | yes | (*5*) | |
| | c.625C>T | | 6m | no | no | yes | yes | (*6*) | |
| | c.625C>T | | birth | yes | yes | yes | yes | (*44*) | |
| | c.625C>T | | 7m | yes | yes | yes | no | (*44*) | |
| C215Y | c.644G>A | 3 | 12y | no | yes | no | no | (*61*) | NEDIM |
| | | | 5y | no | yes | no | no | (*62*) | 2400 ± 982 (n=3): |
| | | | 3y | no | yes | no | no | (*62*) | Very late |
| A227V | c.680C>T | 2 | 2m | yes | yes | yes | yes | (*50*) | DEE17 |
| | | | birth | yes | yes | yes | no | (*63*) | 31 ± 30 (n=2): Early |
| Y231C | c.692A>G | 3 | 5d | yes | no | yes | yes | (*6*) | DEE17 |
| | | | 3d | yes | yes | yes | no | (*46*) | 23 ± 19 (n=3): Early |
| | c.692A>G | | 1m | yes | yes | yes | yes | (*64*) | |
| Q233P | c.698A>C | 1 | 2y | no | yes | no | no | (*65*) | NEDIM<br>720 (n=1): Very late |
| E237K | c.709G>A | 8 | 6m | no | yes | yes | yes | (*5*) | NEDIM |
| | c.709G>A | | 3m | no | yes | yes | yes | (*56*) | 666 ± 316 (n=5): Late |
| | c.709G>A | | infancy | no | yes | yes | yes | (*60*) | |
| | c.709G>A | | infancy | no | yes | yes | no | (*60*) | |
| | c.709G>A | | 6m | no | yes | yes | N/A | (*66*) | |
| | c.709G>A | | 4y | no | yes | yes | no | (*67*) | |
| | c.709G>A | | 4y | no | yes | yes | N/A | (*67*) | |
| | c.709G>A | | N/A | no | yes | yes | N/A | (*68*) | |
| E246K | c.736G>A | 13 | 4m | no | yes | yes | no | (*50*) | NEDIM |
| | | | 11m | no | yes | yes | no | (*52*) | 278 ± 109 (n=12): Late |
| | | | 5m | yes | yes | yes | yes | (*52*) | |
| | c.736G>A | | 3m | no | yes | yes | yes | (*56*) | |
| | | | 3m | no | yes | yes | yes | (*63*) | |
| | c.736G>A | | 3m | no | yes | yes | no | (*69*) | |
| | c.736G>A | | 3m | no | yes | yes | yes | (*69*) | |
| | c.736G>A | | 6m | no | yes | yes | yes | (*69*) | |
| | c.736G>A | | 5m | no | yes | yes | yes | (*69*) | |
| | | | 9m | no | yes | yes | yes | (*70*) | |

| | | | | | | | | | |
|---|---|---|---|---|---|---|---|---|---|
| | c.736G>A<br>c.736G>A<br>c.736G>A | | 11m<br>4y<br>childhood | no<br>no<br>yes | yes<br>yes<br>N/A | yes<br>yes<br>N/A | no<br>yes<br>N/A | (*71*)<br>(*71*)<br>(*72*) | |
| N270H | c.808A>C | 1 | 3m | yes | yes | yes | yes | (*73*) | DEE17<br>90 (n=1): Early |
| F275S | c.824T>C | 1 | 3d | yes | yes | no | yes | (*73*) | DEE17<br>3 (n=1): Very early |
| I279N | c.836T>A<br>c.836T>A<br>c.836T>A | 3 | 4d<br>1h<br>26min | yes<br>yes<br>yes | no<br>yes<br>yes | yes<br>yes<br>yes | yes<br>yes<br>yes | (*4*)<br>(*6*)<br>(*74*) | DEE17<br>2 ± 1 (n=3): Very early |

**Table S2. Oligonucleotide primers used in this study.**

| Name | Sequence | Usage |
|---|---|---|
| Gαo-40-For | 5′-TGCTCAGGGCTGGAGAATCAGG-3′ | Generation of Gαo variants |
| Gao-40-Rev | 5′-CCTGATTCTCCAGCCCTGAGCA-3′ | |
| Gαo-45-For | 5′-CTGGAGAATCAGAAAAAAGCACCATT-3′ | |
| Gao-45-Rev | 5′-AATGGTGCTTTTTTCTGATTCTCCAG-3′ | |
| Gαo-47-For | 5′-CTGGAGAATCAGGAAAAGGCACCATT-3′ | |
| Gao-47-Rev | 5′-AATGGTGCCTTTTCCTGATTCTCCAG-3′ | |
| Gαo-174-For | 5′-AGCAGGGCATCCTCCGAACCAG-3′ | |
| Gao-174-Rev | 5′-CTGGTTCGGAGGATGCCCTGCT-3′ | |
| Gαo-199-For | 5′-AGAACCTCCACTTCAGGCCGTTTG-3′ | |
| Gao-199-Rev | 5′-CAAACGGCCTGAAGTGGAGGTTCT-3′ | |
| Gαo-227-For | 5′-TGTGTCGTGCTCAGCGGCTATGACCAGGTGCT-3′ | |
| Gao-227-Rev | 5′-AGCACCTGGTCATAGCCGCTGAGCACGACACA-3′ | |
| Gαo-231-For | 5′-TGTGTCGCGCTCAGCGGCTGTGACCAGGTGCT-3′ | |
| Gao-231-Rev | 5′-AGCACCTGGTCACAGCCGCTGAGCGCGACACA-3′ | |
| Gαo-233-For | 5′-TGTGTCGCGCTCAGCGGCTATGACCCGGTGCT-3′ | |
| Gao-233-Rev | 5′-AGCACCGGGTCATAGCCGCTGAGCGCGACACA-3′ | |
| Gαo-270-For | 5′-TTCCTCCACAAGAAAGATCTCTTTGGCGAGAA-3′ | |
| Gao-270-Rev | 5′-TTCTCGCCAAAGAGATCTTTCTTGTGGAGGAA-3′ | |
| Gαo-275-For | 5′-TTCCTCAACAAGAAAGATCTCTCTGGCGAGAA-3′ | |
| Gao-275-Rev | 5′-TTCTCGCCAGAGAGATCTTTCTTGTTGAGGAA-3′ | |
| Gαo-279-For | 5′-TGGCGAGAAGAACAAGAAGTCACCT-3′ | |
| Gao-279-Rev | 5′-AGGTGACTTCTTGTTCTTCTCGCCA-3′ | |
| dGαo-203-For | 5′-TTACGTTCCGAGCGCTGACCGCGCACGTCAAACAATTTAA-3′ | Generation of *Drosophila* Gao G203R |
| dGao-203-Rev | 5′-TTAAATTGTTTGACGTGCGCGGTCAGCGCTCGGAACGTAA-3′ | |
| Ric8A-SalI-For | 5′-GCGTCGtCTTCgtcgacCCGGTGCCAGGGGCCATG-3′ | Generation of GFP-Ric8A wild-type and mutants |
| Ric8A-PspOMI-Rev | 5′-GGGAGCAgGGCCcCTGGCATCTTCAGTCAGGATCT-3′ | |
| Ric8A-R75E-For | 5′-CTATCCGAATCCTATCCgaAGACCGCAGCTGCCTGG-3′ | |
| Ric8A-R75E-Rev | 5′-CCAGGCAGCTGCGGTCTtcGGATAGGATTCGGATAG-3′ | |
| Ric8A-K225A-For | 5′-GTGATATTAAAGAGCACTgcGAGGATCTCCATGGCC-3′ | |
| Ric8A-K225A-Rev | 5′-GGCCATGGAGATCCTCgcAGTGCTCTTTAATATCAC-3′ | |
| Ric8B-XhoI-For | 5′-agcctgagctcgtttCTCgaGcggccgccaccatggatga-3′ | Generation of GFP-Ric8B |
| Ric8B-EcoRI-Rev | 5′-cctgtgtggcgaattctcagtctgtgtccgagctg-3′ | |
| dRic8-BsrGI-For | 5′-ATACAAGTTTGTACAAACAAGCAGGCTCGAGGGCCGCCGCCTTCACCATG-3′ | Generation of *Drosophila* GFP-Ric8 |
| dRic8-PspOMI-Rev | 5′-CTGGGTCGGCGGGCCCACCCTAGGTTTTCCGCTTC-3′ | |

**References**


1. W. M. Oldham, H. E. Hamm, Heterotrimeric G protein activation by G-protein-coupled receptors. *Nat Rev Mol Cell Biol* **9**, 60-71 (2008).
2. E. M. Ross, T. M. Wilkie, GTPase-activating proteins for heterotrimeric G proteins: regulators of G protein signaling (RGS) and RGS-like proteins. *Annu Rev Biochem* **69**, 795-827 (2000).
3. C. Lin *et al.*, Double suppression of the Galpha protein activity by RGS proteins. *Molecular cell* **53**, 663-671 (2014).
4. K. Nakamura *et al.*, De Novo mutations in GNAO1, encoding a Galphao subunit of heterotrimeric G proteins, cause epileptic encephalopathy. *Am J Hum Genet* **93**, 496-505 (2013).
5. T. Schirinzi *et al.*, Phenomenology and clinical course of movement disorder in GNAO1 variants: Results from an analytical review. *Parkinsonism Relat Disord* **61**, 19-25 (2019).
6. M. Kelly *et al.*, Spectrum of neurodevelopmental disease associated with the GNAO1 guanosine triphosphate-binding region. *Epilepsia* **60**, 406-418 (2019).
7. E. Axeen *et al.*, Results of the First GNAO1-Related Neurodevelopmental Disorders Caregiver Survey. *Pediatr Neurol* **121**, 28-32 (2021).
8. M. Savitsky, G. P. Solis, M. Kryuchkov, V. L. Katanaev, Humanization of Drosophila Gαo to Model GNAO1 Paediatric Encephalopathies. *Biomedicines* **8**, 395 (2020).
9. B. S. Muntean *et al.*, Gαo is a major determinant of cAMP signaling in the pathophysiology of movement disorders. *Cell Rep* **34**, 108718 (2021).
10. Y. A. Larasati *et al.*, Restoration of the GTPase activity and cellular interactions of Galpha(o) mutants by Zn(2+) in GNAO1 encephalopathy models. *Sci Adv* **8**, eabn9350 (2022).
11. D. Wang *et al.*, Genetic modeling of GNAO1 disorder delineates mechanisms of Gαo dysfunction. *Hum Mol Genet* **31**, 510-522 (2022).
12. H. Feng *et al.*, Movement disorder in GNAO1 encephalopathy associated with gain-of-function mutations. *Neurology* **89**, 762-770 (2017).
13. L. Song *et al.*, Identification of functional cooperative mutations of GNAO1 in human acute lymphoblastic leukemia. *Blood* **137**, 1181-1191 (2021).
14. M. Di Rocco *et al.*, Caenorhabditis elegans provides an efficient drug screening platform for GNAO1-related disorders and highlights the potential role of caffeine in controlling dyskinesia. *Hum Mol Genet* **31**, 929-941 (2022).
15. D. Silachev *et al.*, Mouse models characterize GNAO1 encephalopathy as a neurodevelopmental disorder leading to motor anomalies: from a severe G203R to a milder C215Y mutation. *Acta Neuropathol Commun* **10**, 9 (2022).
16. M. Gabay *et al.*, Ric-8 proteins are molecular chaperones that direct nascent G protein α subunit membrane association. *Sci Signal* **4**, ra79-ra79 (2011).
17. G. P. Solis *et al.*, Pediatric Encephalopathy: Clinical, Biochemical and Cellular Insights into the Role of Gln52 of GNAO1 and GNAI1 for the Dominant Disease. *Cells* **10**, 2749 (2021).
18. C. L. Larrivee *et al.*, Mice with GNAO1 R209H Movement Disorder Variant Display Hyperlocomotion Alleviated by Risperidone. *J Pharmacol Exp Ther* **373**, 24-33 (2020).
19. D. Kopein, V. L. Katanaev, Drosophila GoLoco-protein pins is a target of Galpha(o)-mediated G protein-coupled receptor signaling. *Mol Biol Cell* **20**, 3865-3877 (2009).
20. G. P. Solis *et al.*, Golgi-Resident Galphao Promotes Protrusive Membrane Dynamics. *Cell* **170**, 939-955 (2017).
21. V. L. Katanaev, M. Chornomorets, Kinetic diversity in G-protein-coupled receptor signalling. *Biochem J* **401**, 485-495 (2007).
22. G. P. Solis *et al.*, Local and substrate-specific S-palmitoylation determines subcellular localization of Gαo. *Nat Commun* **13**, 2072 (2022).
23. G. P. Solis, V. L. Katanaev, Galphao (GNAO1) encephalopathies: plasma membrane vs. Golgi functions. *Oncotarget* **9**, 23846-23847 (2018).
24. H. Schihada, R. Shekhani, G. Schulte, Quantitative assessment of constitutive G protein-coupled receptor activity with BRET-based G protein biosensors. *Sci Signal* **14**, eabf1653 (2021).
25. G. G. Tall, A. M. Krumins, A. G. Gilman, Mammalian Ric-8A (synembryn) is a heterotrimeric Galpha protein guanine nucleotide exchange factor. *Journal of Biological Chemistry* **278**, 8356-8362 (2003).

26. A. B. Seven *et al.*, Structures of Gα Proteins in Complex with Their Chaperone Reveal Quality Control Mechanisms. *Cell Rep* **30**, 3699-3709.e3696 (2020).
27. H. J. Muller, Further studies on the nature and causes of gene mutations. *Proceedings of the Sixth International Congress of Genetics, Ithaca, New York.* **1**, 213-255 (1932).
28. V. Takiar, C. K. Ip, M. Gao, G. B. Mills, L. W. Cheung, Neomorphic mutations create therapeutic challenges in cancer. *Oncogene* **36**, 1607-1618 (2017).
29. P. Chan, M. Gabay, F. A. Wright, G. G. Tall, Ric-8B is a GTP-dependent G protein alphas guanine nucleotide exchange factor. *J Biol Chem* **286**, 19932-19942 (2011).
30. M. M. Papasergi-Scott *et al.*, Structures of Ric-8B in complex with Galpha protein folding clients reveal isoform specificity mechanisms. *Structure*, (2023).
31. A. Koval *et al.*, In-depth molecular profiling of an intronic GNAO1 mutant as the basis for personalized high-throughput drug screening. *Med* **4**, in press (2023).
32. T. Wirth *et al.*, Highlighting the Dystonic Phenotype Related to GNAO1. *Mov Disord* **37**, 1547-1554 (2022).
33. A. M. Luchtenborg *et al.*, Heterotrimeric Go protein links Wnt-Frizzled signaling with ankyrins to regulate the neuronal microtubule cytoskeleton. *Development* **141**, 3399-3409 (2014).
34. G. P. Solis *et al.*, Golgi-Resident Galphao Promotes Protrusive Membrane Dynamics. *Cell* **170**, 939-955 e924 (2017).
35. G. P. Solis *et al.*, Pediatric Encephalopathy: Clinical, Biochemical and Cellular Insights into the Role of Gln52 of GNAO1 and GNAI1 for the Dominant Disease. *Cells* **10**, (2021).
36. S. C. Wang *et al.*, Regulation of type V adenylate cyclase by Ric8a, a guanine nucleotide exchange factor. *Biochem J* **406**, 383-388 (2007).
37. E. M. Jones *et al.*, A Scalable, Multiplexed Assay for Decoding GPCR-Ligand Interactions with RNA Sequencing. *Cell Syst* **8**, 254-260 e256 (2019).
38. K. A. Peters, S. L. Rogers, Drosophila Ric-8 interacts with the Galpha12/13 subunit, Concertina, during activation of the Folded gastrulation pathway. *Mol Biol Cell* **24**, 3460-3471 (2013).
39. G. P. Solis *et al.*, Local and substrate-specific S-palmitoylation determines subcellular localization of Galphao. *Nat Commun* **13**, 2072 (2022).
40. Y. Katoh, S. Nozaki, D. Hartanto, R. Miyano, K. Nakayama, Architectures of multisubunit complexes revealed by a visible immunoprecipitation assay using fluorescent fusion proteins. *J Cell Sci* **128**, 2351-2362 (2015).
41. F. Madeira *et al.*, Search and sequence analysis tools services from EMBL-EBI in 2022. *Nucleic Acids Res* **50**, W276-279 (2022).
42. A. M. Waterhouse, J. B. Procter, D. M. Martin, M. Clamp, G. J. Barton, Jalview Version 2--a multiple sequence alignment editor and analysis workbench. *Bioinformatics* **25**, 1189-1191 (2009).
43. C. Y. Law *et al.*, Clinical whole-exome sequencing reveals a novel missense pathogenic variant of GNAO1 in a patient with infantile-onset epilepsy. *Clin Chim Acta* **451**, 292-296 (2015).
44. F. R. Danti *et al.*, GNAO1 encephalopathy: Broadening the phenotype and evaluating treatment and outcome. *Neurol Genet* **3**, e143 (2017).
45. T. U. J. Bruun *et al.*, Prospective cohort study for identification of underlying genetic causes in neonatal encephalopathy using whole-exome sequencing. *Genet Med* **20**, 486-494 (2018).
46. X. Yang *et al.*, Phenotypes of GNAO1 Variants in a Chinese Cohort. *Frontiers in Neurology* **12**, (2021).
47. P. Gawlinski *et al.*, PEHO Syndrome May Represent Phenotypic Expansion at the Severe End of the Early-Onset Encephalopathies. *Pediatric Neurology* **60**, 83-87 (2016).
48. A. Marcé-Grau *et al.*, GNAO1 encephalopathy: further delineation of a severe neurodevelopmental syndrome affecting females. *Orphanet Journal of Rare Diseases* **11**, 38 (2016).
49. J. Lee *et al.*, Genomic Analysis of Korean Patient With Microcephaly. *Front Genet* **11**, 543528-543528 (2021).
50. H. Saitsu *et al.*, Phenotypic spectrum of GNAO1 variants: epileptic encephalopathy to involuntary movements with severe developmental delay. *Eur J Hum Genet* **24**, 129-134 (2016).
51. R. Arya, C. Spaeth, D. L. Gilbert, J. L. Leach, K. D. Holland, GNAO1-associated epileptic encephalopathy and movement disorders: c.607G>A variant represents a probable mutation hotspot with a distinct phenotype. *Epileptic Disord* **19**, 67-75 (2017).

52. D. C. Schorling *et al.*, Expanding Phenotype of De Novo Mutations in GNAO1: Four New Cases and Review of Literature. *Neuropediatrics* **48**, 371-377 (2017).
53. M. J. Malaquias *et al.*, GNAO1 mutation presenting as dyskinetic cerebral palsy. *Neurol Sci* **40**, 2213-2216 (2019).
54. M. Akasaka *et al.*, GNAO1 mutation-related severe involuntary movements treated with gabapentin. *Brain Dev* **43**, 576-579 (2021).
55. A. K. Kwong *et al.*, Exome sequencing in paediatric patients with movement disorders. *Orphanet J Rare Dis* **16**, 32 (2021).
56. M. Waak *et al.*, GNAO1-related movement disorder with life-threatening exacerbations: movement phenomenology and response to DBS. *J Neurol Neurosurg Psychiatry* **89**, 221-222 (2018).
57. I. Dzinovic *et al.*, Dystonia as a prominent presenting feature in developmental and epileptic encephalopathies: A case series. *Parkinsonism Relat Disord* **90**, 73-78 (2021).
58. P. Danhofer *et al.*, Brittle Biballism-Dystonia in a Pediatric Patient with GNAO1 Mutation Managed Using Pallidal Deep Brain Stimulation. *Mov Disord Clin Pract* **8**, 153-155 (2021).
59. M. Chopra *et al.*, Mendelian etiologies identified with whole exome sequencing in cerebral palsy. *Annals of Clinical and Translational Neurology* **9**, 193-205 (2022).
60. A. Koy *et al.*, Deep brain stimulation is effective in pediatric patients with GNAO1 associated severe hyperkinesia. *J Neurol Sci* **391**, 31-39 (2018).
61. M. Carecchio *et al.*, Frequency and phenotypic spectrum of KMT2B dystonia in childhood: A single-center cohort study. *Movement Disorders* **34**, 1516-1527 (2019).
62. T. Wirth *et al.*, Increased diagnostic yield in complex dystonia through exome sequencing. *Parkinsonism Relat Disord* **74**, 50-56 (2020).
63. S. Y. Kim *et al.*, Spectrum of movement disorders in GNAO1 encephalopathy: in-depth phenotyping and case-by-case analysis. *Orphanet J Rare Dis* **15**, 343 (2020).
64. I. Talvik *et al.*, Clinical Phenotype of De Novo GNAO1 Mutation:Case Report and Review of Literature. *Child Neurol Open* **2**, 2329048X15583717 (2015).
65. S. Yilmaz *et al.*, Excellent response to deep brain stimulation in a young girl with GNAO1-related progressive choreoathetosis. *Childs Nerv Syst* **32**, 1567-1568 (2016).
66. A. Okumura *et al.*, A patient with a GNAO1 mutation with decreased spontaneous movements, hypotonia, and dystonic features. *Brain Dev* **40**, 926-930 (2018).
67. Z. Al Masseri, M. AlSayed, Gonadal mosaicism in GNAO1 causing neurodevelopmental disorder with involuntary movements; two additional variants. *Molecular Genetics and Metabolism Reports* **31**, 100864 (2022).
68. E. L. Fung *et al.*, Deep brain stimulation in a young child with GNAO1 mutation - Feasible and helpful. *Surg Neurol Int* **13**, 285 (2022).
69. A. L. Ananth *et al.*, Clinical Course of Six Children With GNAO1 Mutations Causing a Severe and Distinctive Movement Disorder. *Pediatr Neurol* **59**, 81-84 (2016).
70. Y. Takezawa *et al.*, Genomic analysis identifies masqueraders of full-term cerebral palsy. *Ann Clin Transl Neurol* **5**, 538-551 (2018).
71. A. Benato *et al.*, Long-term effect of subthalamic and pallidal deep brain stimulation for status dystonicus in children with methylmalonic acidemia and GNAO1 mutation. *J Neural Transm (Vienna)* **126**, 739-757 (2019).
72. K. L. Helbig *et al.*, Diagnostic exome sequencing provides a molecular diagnosis for a significant proportion of patients with epilepsy. *Genet Med* **18**, 898-905 (2016).
73. De novo mutations in synaptic transmission genes including DNM1 cause epileptic encephalopathies. *Am J Hum Genet* **95**, 360-370 (2014).
74. De Novo Mutations in SLC1A2 and CACNA1A Are Important Causes of Epileptic Encephalopathies. *Am J Hum Genet* **99**, 287-298 (2016).

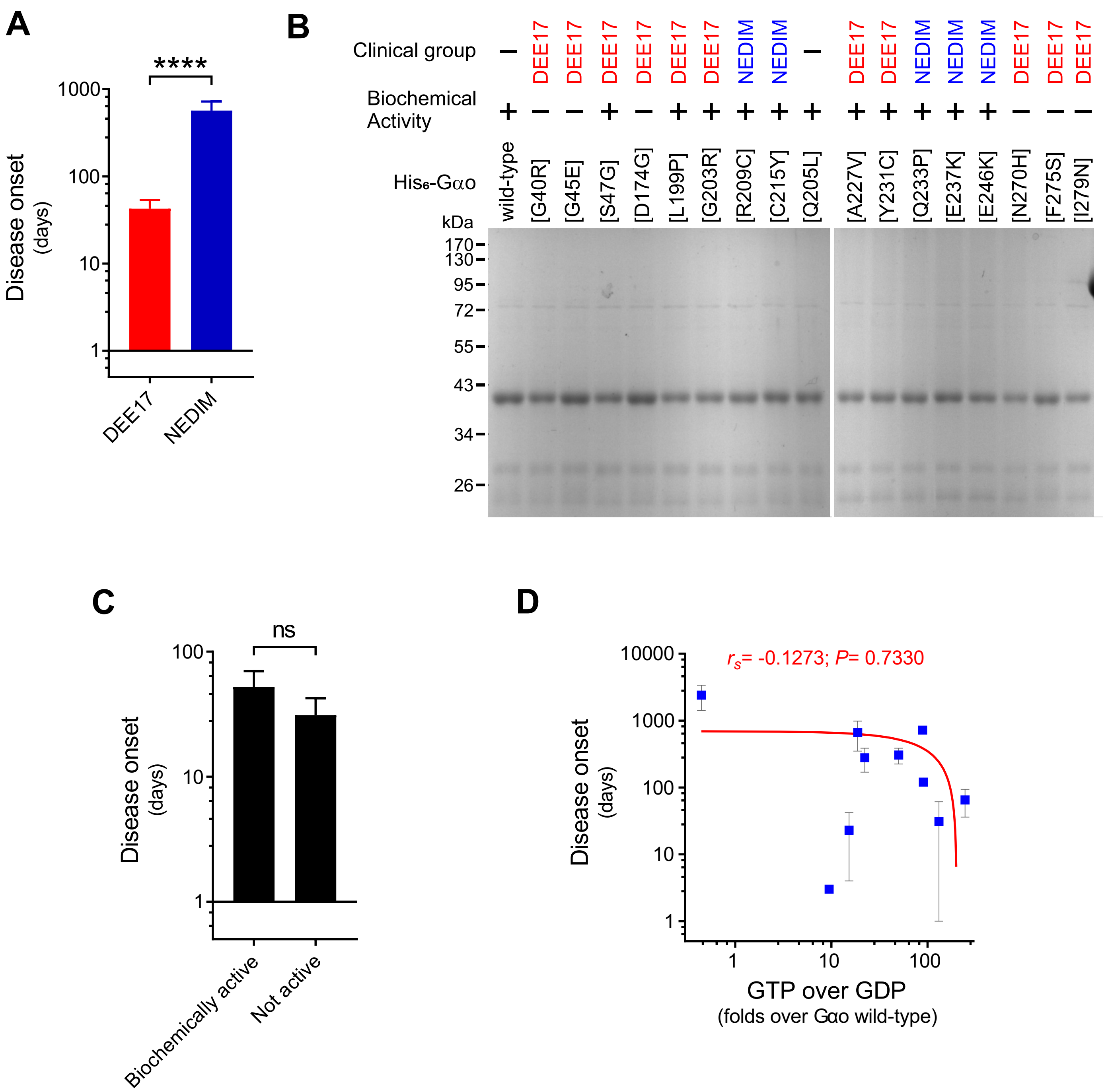


**Fig. S1. Biochemical purification of Gαo encephalopathy mutants.** **(A)** The Disease onset data from patients **(Table S1)** was pooled according to the two classifications of *GNAO1* encephalopathy: Developmental and Epileptic Encephalopathy-17 (DEE17; red bars) and Neurodevelopmental Disorder with Involuntary Movements (NEDIM; blue bars). Note that the most severe DEE17 group shows in average a much lower Disease onset than the NEDIM group ($n$=29-31). **(B)** Coomasie blue staining of SDS-PAGE shows the purity of the recombinant $His_6$-tagged Gαo wild-type, encephalopathy mutants, and the control Q205L. The clinical manifestation associated to each Gαo mutant, and if they were purified active (+) or not active (-) is indicated. **(C)** The Disease onset data was grouped according to the biochemical activity of the recombinant Gαo mutants associated to the DEE17 disorder ($n$=13-16). **(D)** A scatterplot shows a non-significant negative correlation between Disease onset and the calculated GTP/GDP-loading ratio of Gαo variants. Note the log scale in the *y* axis. Data shown as means ± SEM. Data in **(A)** and **(C)** were analyzed by two-tailed Mann Whitney test, and in **(D)** by two-tailed Spearman correlation test (rank correlation coefficient ($r_s$) and $P$ value are indicated). ns is not significant and ****$P$ < 0.0001.

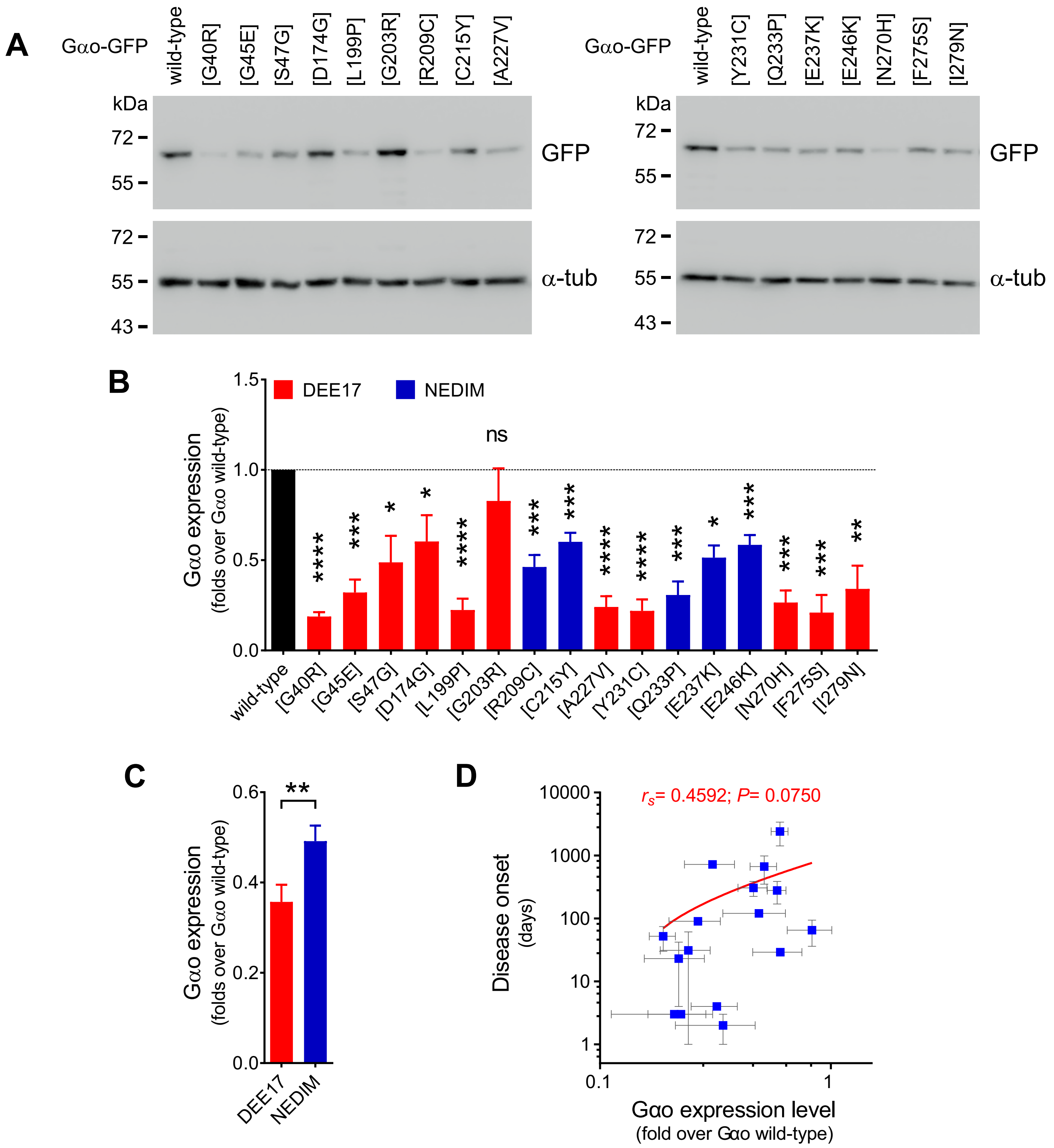


**Fig. S2. Expression of Gαo encephalopathy mutants in N2a cells. (A)** N2a cells were transfected with Gαo-GFP wild-type or encephalopathy mutants, and their expression levels were determined by Western blot using antibodies against GFP and against α-tubulin (α-tub) as loading control. **(B)** Quantification of the expression levels of Gαo variants ($n$=6). Data are color-coded according to the involvement of the mutants in Developmental and Epileptic Encephalopathy-17 (DEE17; red bars) or Neurodevelopmental Disorder with Involuntary Movements (NEDIM; blue bars). **(C)** The expression level of Gαo mutants pooled according to the DEE17 and NEDIM classification ($n$=27-66). **(D)** A scatterplot shows no significant correlation between Disease onset and the expression of Gαo variants. Note the log scale in the *y* axis. All data are shown as means ± SEM. Data in **(B)** were analyzed by one-sample t-test, **(C)** by two-tailed Mann Whitney test, and **(D)** by two-tailed Spearman correlation test (rank correlation coefficient ($r_s$) and *P* value are indicated). ns is not significant, $*P < 0.05$, $**P < 0.01$, $***P < 0.001$ and $****P < 0.0001$.

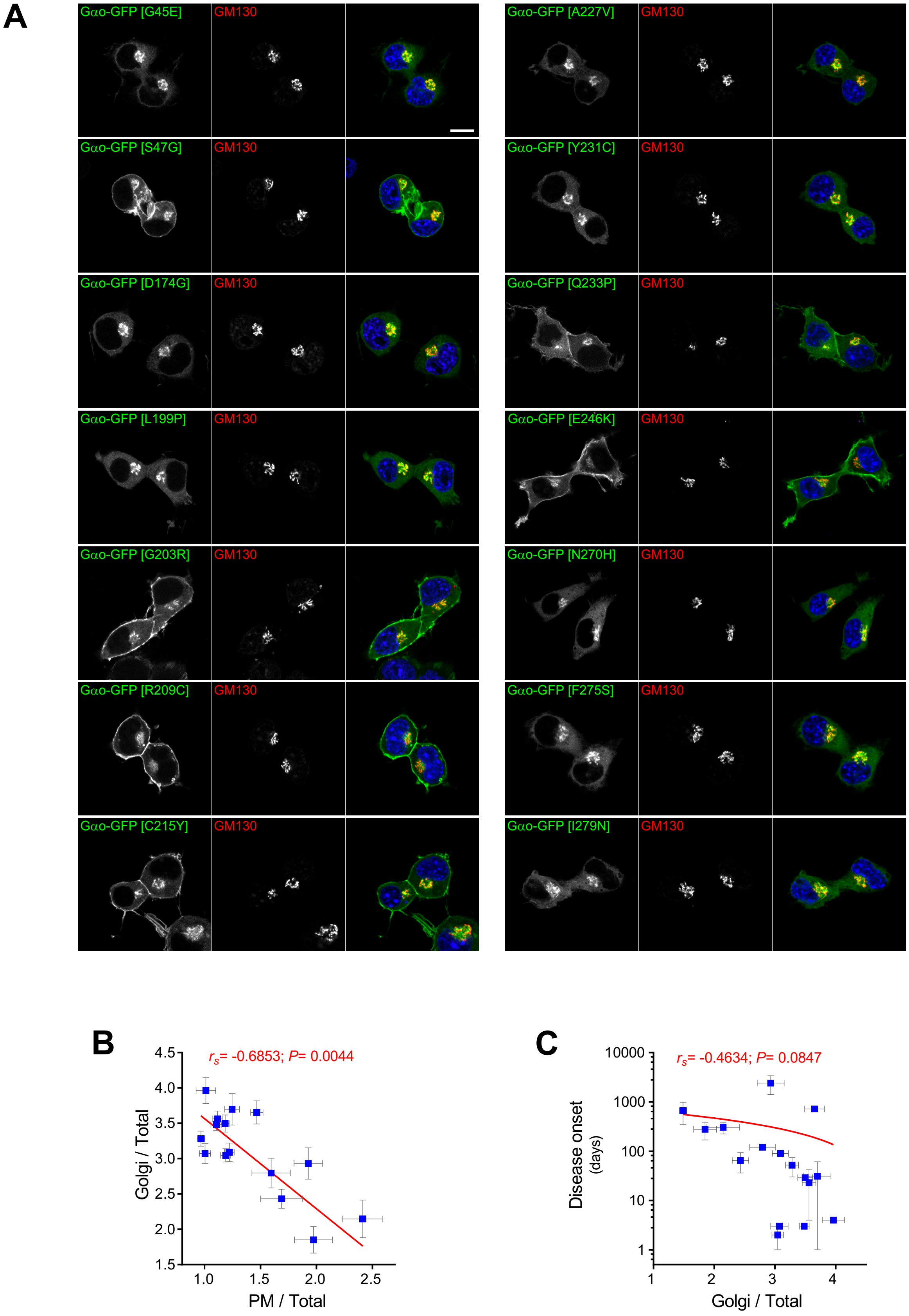


**Fig. S3. Subcellular localization of Gαo encephalopathy mutants in N2a cells. (A)** N2a cells expressing Gαo-GFP wild-type or the indicated encephalopathy mutants were immunostained against GM130 to visualize the Golgi apparatus. Scale bar, 10 µm. **(B)** A scatterplot shows a significant negative correlation between the relative localization of Gαo mutants at the plasma membrane (PM) and Golgi apparatus. **(C)** No significant correlation was calculated between Disease onset and the Golgi localization of Gαo variants. Note the log scale in the *y* axis. Data shown as means ± SEM. Data in **(B)** and **(C)** were analyzed by two-tailed Spearman correlation test; rank correlation coefficients ($r_s$) and *P* values are indicated.

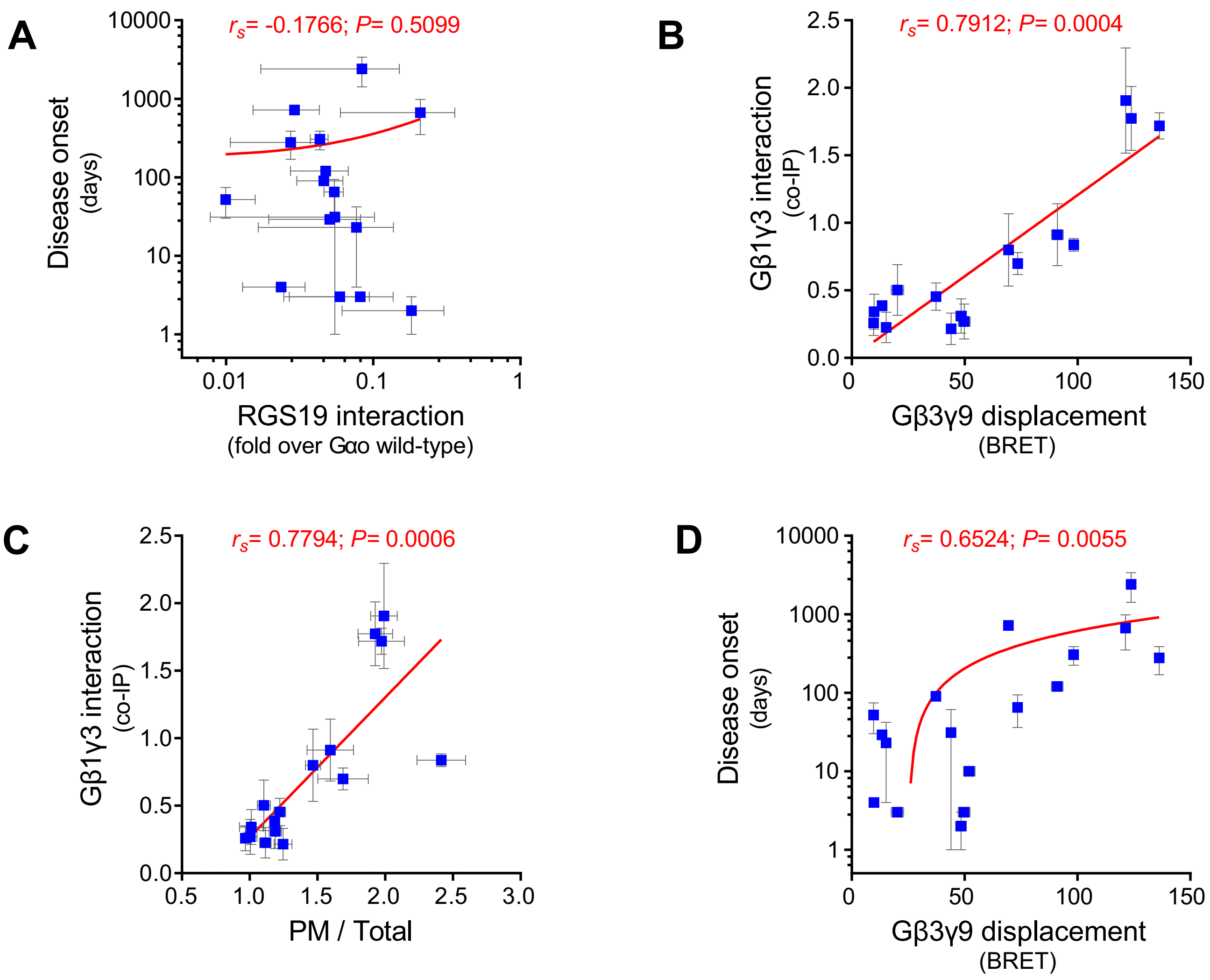


**Fig. S4. Analysis of the cellular properties of Gαo encephalopathy mutants. (A)** A scatterplot shows no significant correlation between Disease onset and RGS19 interaction of Gαo variants. Note the log scale in the *y* axis. **(B and C)** Strong positive correlations were calculated between Gβ1γ3 interaction and Gβ3γ9 displacement **(B)** as well as plasma membrane (PM) localization **(C)** of Gαo mutants. **(D)** A significant positive correlation is also seen between Disease onset and Gβ3γ9 displacement. Note the log scale in the *y* axis. Data shown as means ± SEM. All data were analyzed by two-tailed Spearman correlation test; rank correlation coefficients ($r_s$) and $P$ values are indicated.

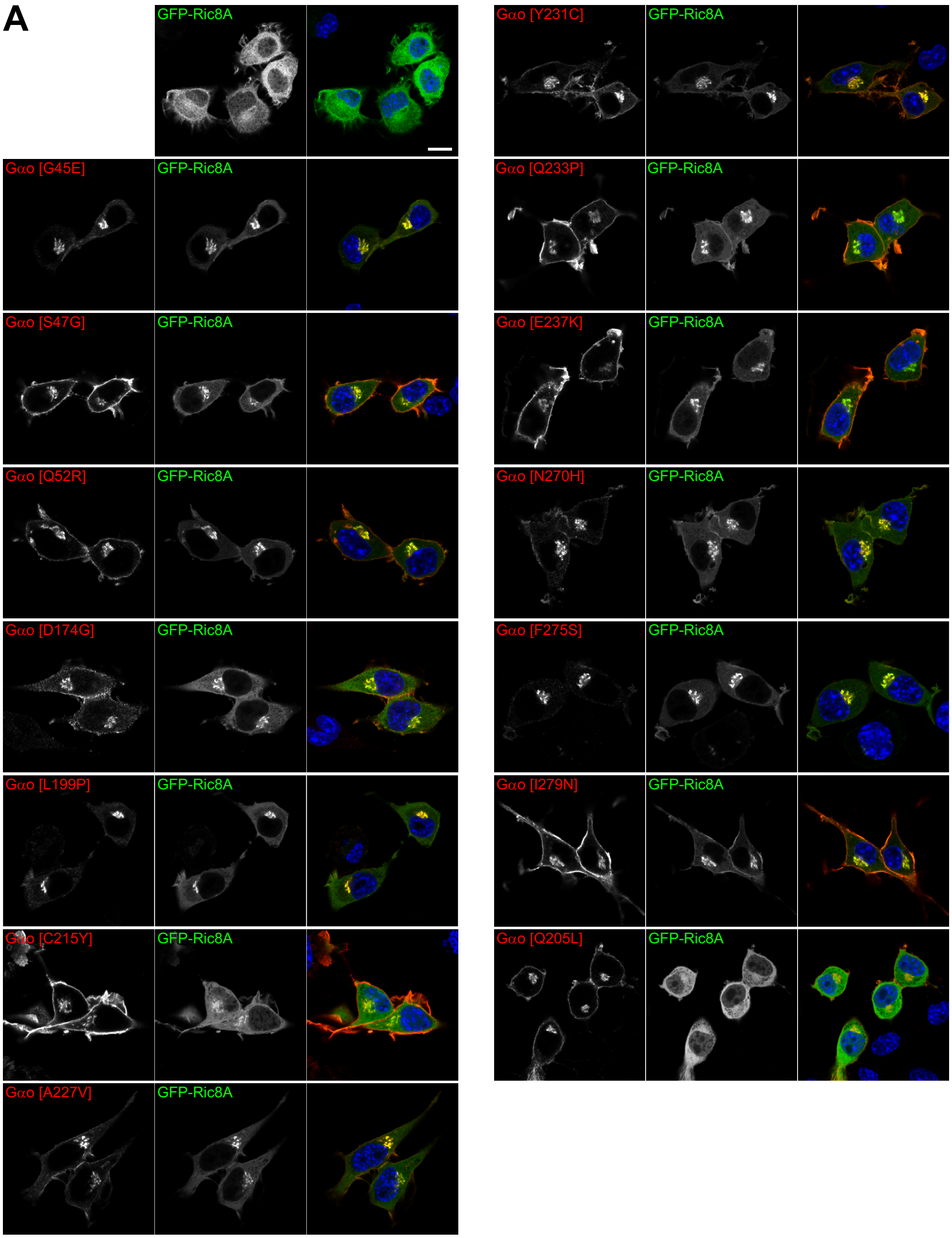


**Fig. S5. Golgi delocalization of Ric8A by Gαo encephalopathy mutants. (A)** Representative images of N2a cells expressing GFP-Ric8A alone, together with Gαo encephalopathy mutants, or the GTPase-deficient Q205L mutant as control. Note that the normal cytoplasmic localization of Ric8A is drastically changed to the Golgi, and to a lesser extent to the plasma membrane, by the co-expression of Gαo encephalopathy variants, but not Q205L. Gαo was detected by immunostaining using an specific antibody. Scale bar, 10 µm.

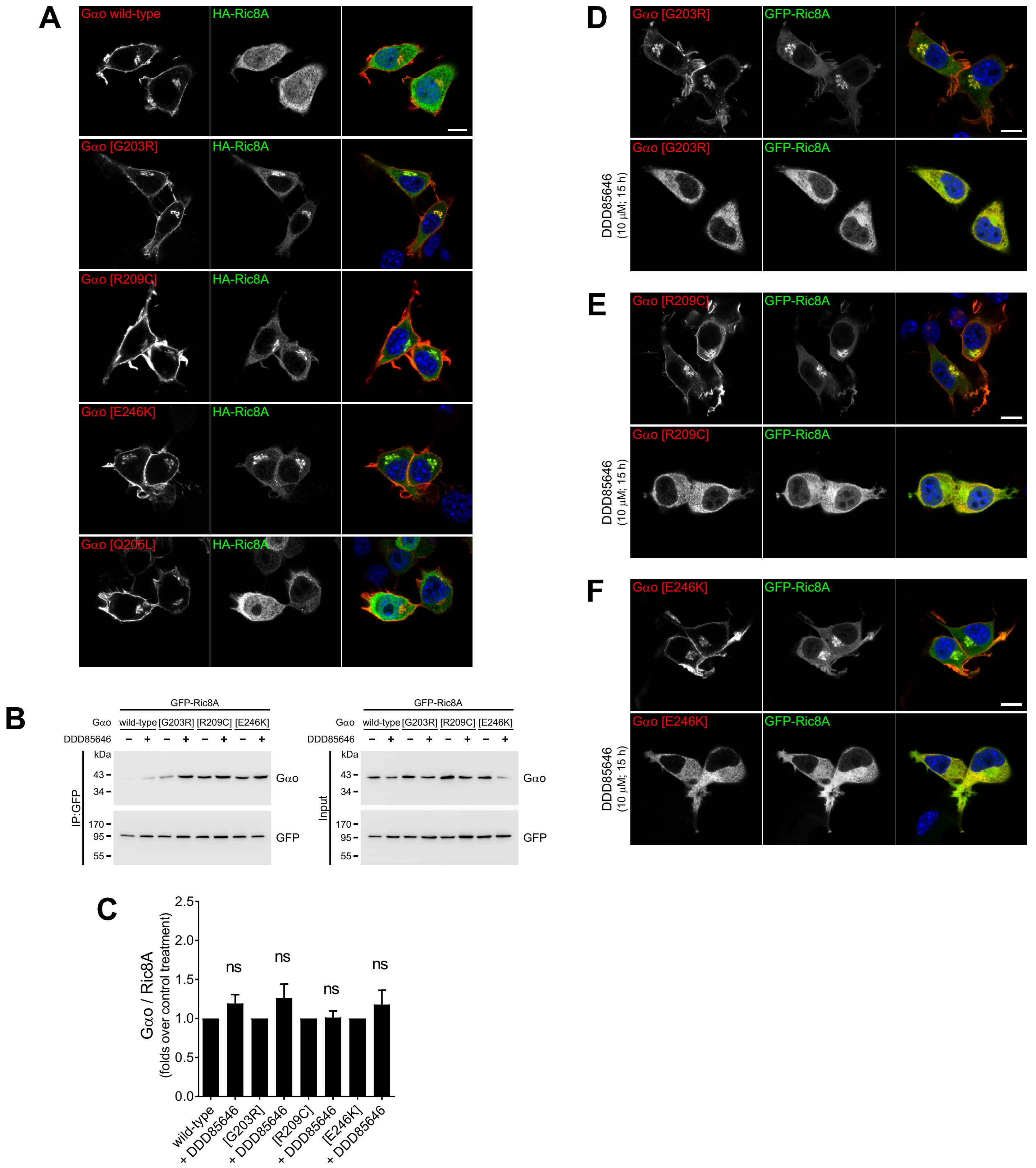


**Fig. S6. Golgi delocalization of Ric8A by encephalopathy mutants depends on Gαo lipidations. (A)** N2a cells co-expressing HA-tagged Ric8A (HA-Ric8A) together with Gαo wild-type, the mutants G203R, R209C, E246K, or the control Q205L were immunostained against Gαo and the HA-epitope. Note the strong cytoplasm-to-Golgi delocalization of Ric8A only in the presence of encephalopathy mutants. **(B)** Immunoprecipitation (IP) of GFP-Ric8A was done from N2a cells preincubated for 15 h with 10 μM of the N-myristoylation blocker DDD85646 (+) or DMSO as control (-), and using a nanobody against GFP. The co-precipitation of Gαo variants was determined by immunodetection with antibodies against Gαo and GFP. **(C)** Quantification of Gαo co-IP reveals no significant effect of the N-myristoylation inhibitor ($n$=3-4). **(D to F)** Representative images of N2a cells showing that the Golgi delocalization of Ric8A and overall membrane association of Gαo variants were abolished by the DDD85646 treatment. Scale bars, 10 μm. Data shown as means ± SEM. The data in **(C)** were analyzed by one-sample t-test; ns is not significant.

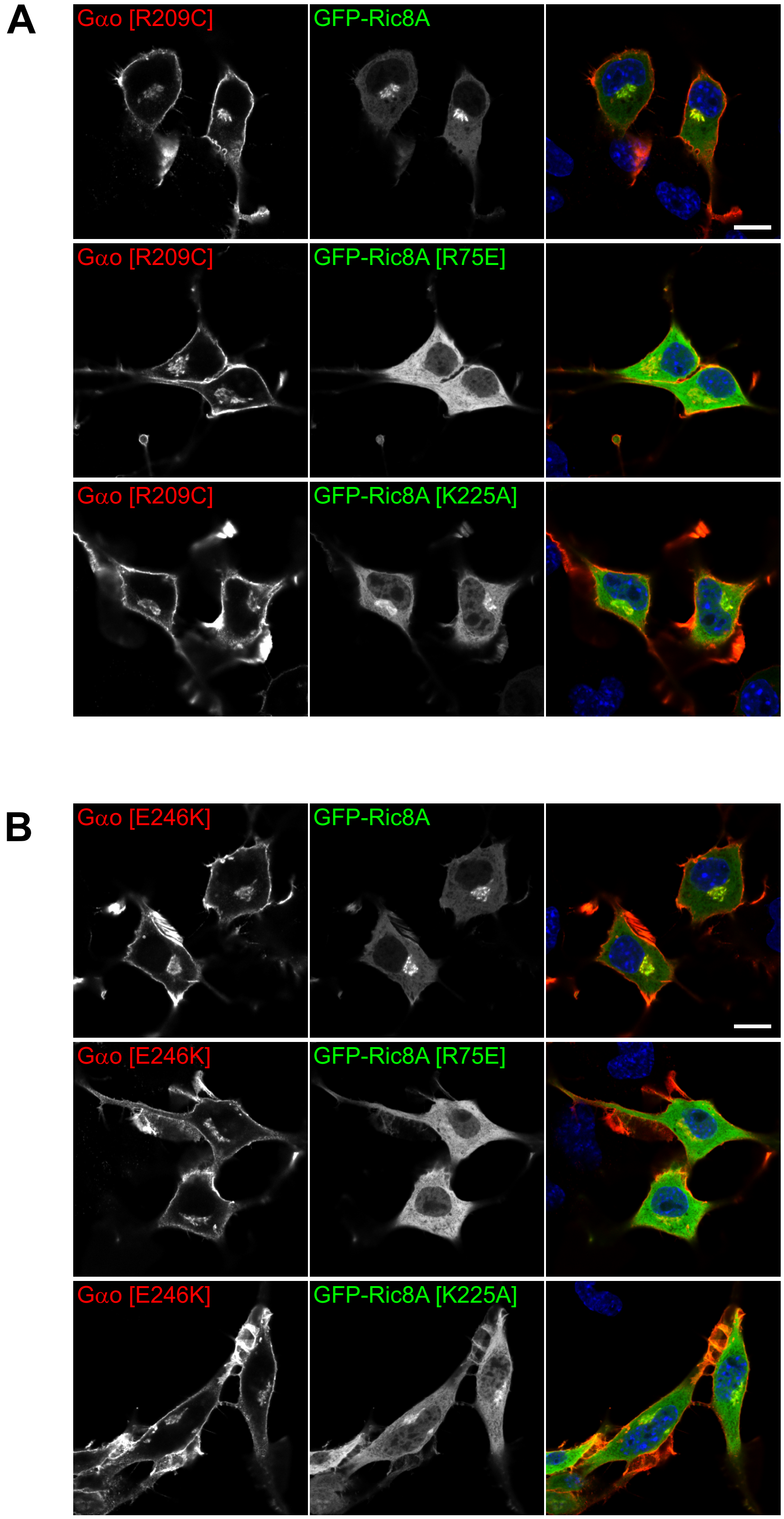


**Fig. S7. The neomorphic Gαo-Ric8A interaction of encephalopathy mutants depends on Ric8A chaperone activity. (A** and **B)** N2a cells co-expressing the GFP-Ric8A and Gαo constructs indicated in the panels were immunostained against Gαo and DAPI staining in blue indicates nuclei. Note that the strong Golgi-delocalization of Ric8A by the Gαo R209C and E246K mutants is clearly reduced or lost for the chaperone-deficient mutants K225A and R75E, respectively. Scale bars, 10 μm.

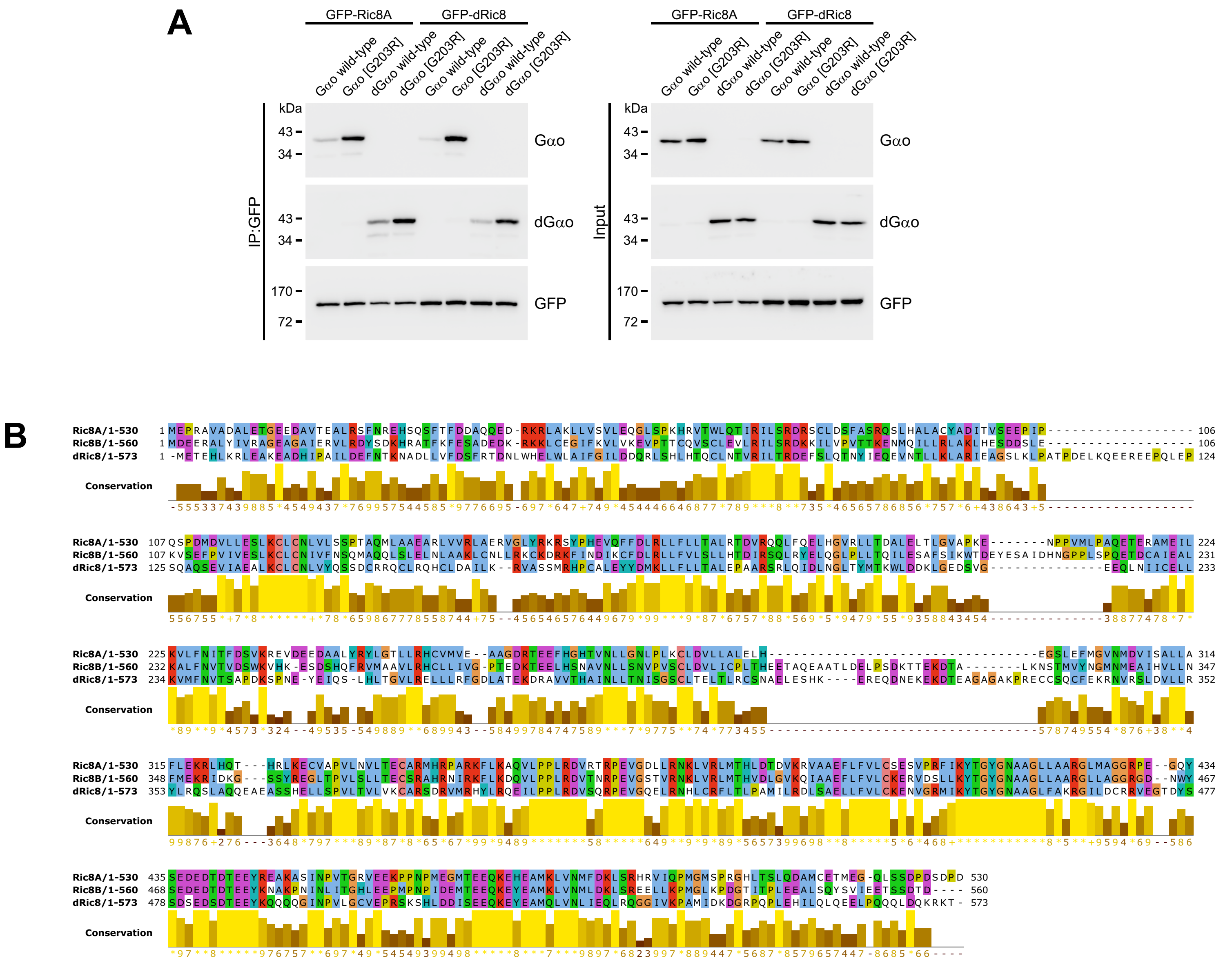


**Fig. S8. The neomorphic Gαo-Ric8 interaction is conserved between fly and mammals. (A)** N2a cells were co-transfected with GFP-Ric8A (mouse) or GFP-dRic8 (*Drosophila*), and the G203R mutant of Gαo (human) and dGαo (*Drosophila*). The immunoprecipitation (IP) of GFP constructs was done with a nanobody against GFP and analyzed by Western blot using antibodies against GFP, Gαo, and dGαo. **(B)** A multiple sequence alignment of Ric8 proteins including Ric8A *Mus musculus* (NP_444424.1), Ric8B *Mus musculus* (NP_898995.1), and dRic8 *Drosophila melanogaster* (NP_001285048.1).
